# Mismatch tolerance of a gRNA for CRISPR-based gene activation confers broad activity critical for cell reprogramming

**DOI:** 10.64898/2026.02.01.703129

**Authors:** Samuel J. Reisman, Wei Zhu, Samantha E. Miller, Dahlia Halabi, Nicholas Sangvai, Gregory E. Crawford, Raluca Gordân, Charles A. Gersbach

## Abstract

CRISPR activation and interference systems (CRISPRa/i) are widely used for programmable transcriptional control. Although these technologies are capable of highly specific single-gene activity, some applications of transcriptional network reprogramming require broad, genome-wide effects. Here, we identify a CRISPRa gRNA that robustly reprograms astrocyte transcriptional state. Unexpectedly, this activity arises from extensive off-target binding that induces expression changes in thousands of genes, unlike neighboring gRNAs targeting the same intended on-target site. We leverage this promiscuous gRNA to dissect determinants of gRNA-driven off-target dCas9 binding in the context of transcriptional reprogramming. Using ChIP-seq, high-throughput protein-binding microarrays, and gRNA-variant library screening in cells, we demonstrate that PAM-proximal bases are primary determinants of genomic binding, mismatch tolerance is both gRNA- and base-specific, and targeted mutations within the PAM-proximal region can tune gRNA specificity. We further demonstrate that CRISPRa-driven phenotypes can reflect combined contributions from widespread off-target activity and dose-dependent on-target effects. These findings highlight the potentially widespread impacts of CRISPRa off-target activity, underscore the need to account for cryptic effects when selecting and evaluating gRNAs for programming cell phenotypes, and demonstrate that multi-site binding by CRISPRa systems can be exploited as a feature for network-level perturbations in cell reprogramming.

**GRAPHICAL ABSTRACT:** 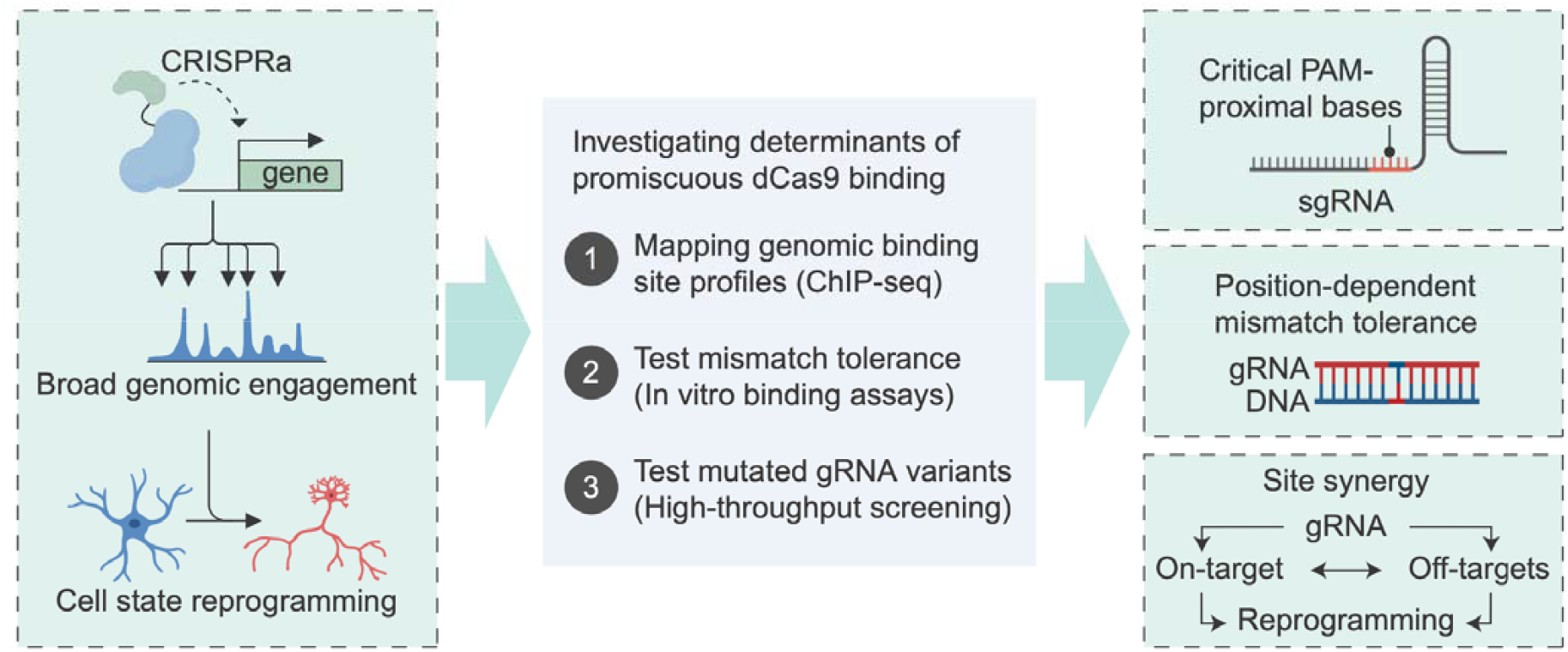

## INTRODUCTION

CRISPR activation and interference systems (CRISPRa/i), commonly consisting of nuclease-deactivated Cas (dCas) protein fused to epigenetic effectors, enable programmable modulation of gene expression with unprecedented ease^1,2^. As a result, they have become increasingly popular tools for high-throughput screening by transcriptional perturbation of genes and non-coding regulatory elements^3,4^ and are increasingly explored in the context of gene therapy^5^, as their lack of DNA cleavage ameliorates concerns of chromosomal rearrangements associated with double-strand breaks (DSBs)^6^. CRISPRa has also been widely used for cell reprogramming applications^7–14^, typically by inducing expression of one or more endogenous genes that encode for master transcription factors (TFs) that coordinate broad gene regulatory networks that define cell state. Thus the primary effect of CRISPRa on the intended target gene can be highly specific, while leading to secondary downstream widespread transcriptional changes. Many studies target known master TFs that have been associated with particular cell phenotypes, but high-throughput CRISPRa phenotypic screens that perturb all TF genes individually and employ readouts including FACS, cell proliferation, and single cell RNA-seq^15–19^ can also be used to discover TFs that coordinate specific cell states.

While it is clear that for each variation of CRISPR technology it is possible to select or engineer gRNAs with exceptional genome-wide specificity, it is also evident that some gRNAs have the potential for variable levels of off-target effects. The lack of DNA cleavage and DNA-level edits from CRISPRa/i presents unique challenges to detecting potential off-target activity. As possible gRNA-driven off-target editing in other CRISPR modalities has become increasingly appreciated, monumental effort has been dedicated to developing tools to map these off-targets, including CIRCLE-seq^20^, GUIDE-seq ^21^, Digenome-seq^22^, and DISCOVER-seq^23^, among others. These tools rely on directly or indirectly sequencing edited DNA and thus cannot be applied to CRISPRa/i. One approach for mapping off-target activity of CRISPRa/i is RNA-sequencing to find modulated transcripts^24^.However, in cases where changes to the direct target gene leads to hundreds or thousands or downstream secondary changes in gene expression, such as when modulating master TFs for cell reprogramming, this strategy is limited in the ability to identify active off-target sites. This is an acute challenge in high-throughput phenotypic CRISPR screens where secondary downstream effects of epigenetic editing are expected. In these contexts, it remains difficult to attribute observed phenotypes solely to intended on-target effects.

Chromatin immunoprecipitation and sequencing (ChIP-seq) and similar technologies can be used for direct mapping of off-target binding of CRISPRa/i systems^25–27^, but remain low-throughput. Previous work employing ChIP-seq has shown that specificity of CRISPRa/i is gRNA-dependent and can range from highly precise to promiscuous^25,28–30^, highlighting the need to better understand the determinants of dCas9 off-target binding. Factors contributing to gRNA promiscuity in CRISPRa/i are not well understood, and few examples exist of functional off-target activity with significant downstream effects. Additionally, across other CRISPR systems, protospacer mismatches are tolerated in a position- and gRNA-dependent manner^31^. However, the interplay of mismatch tolerance, off-target dCas9 binding, and transcriptional changes in the human genome that alter cell state has not been sufficiently explored. This is particularly important in the context of reprogramming where broad transcriptional engagement may itself contribute to functional outcomes.

In this study, we identify a promiscuous CRISPRa gRNA with functional off-target activity in primary astrocytes, leading to altered expression of thousands of genes and reprogramming of astrocyte transcriptional state. We utilize ChIP-seq, high-throughput *in vitro* protein-binding microarrays, and high-throughput screening of gRNA-variant libraries in cells to discern the relationships between PAM-proximal base-driven gRNA binding, protospacer mismatch tolerance, and off-target activity. We report that PAM-proximal gRNA base sequence is a major determinant of off-target binding, that mismatch tolerance varies between gRNAs and base position, and that engineering gRNA variants can alter gRNA promiscuity. Importantly, properties traditionally viewed as undesirable can also enable broad transcriptional network activation that contributes to desired reprogramming outcomes. These findings clarify how gRNA-dependent mismatch tolerance shapes both unintended off-target effects and potentially advantageous broad network engagement, providing a framework for interpreting large-scale CRISPRa/i screens and leveraging these features for transcriptional network rewiring.

## MATERIALS AND METHODS

### Plasmids

Plasmids were constructed using Gibson assembly (NEB). The all-in-one CRISPRa lentiviral plasmid expressing ^VP64^dSpCas9^VP64^, a gRNA scaffold, and a puromycin selection cassette was generated by modifying Addgene plasmid #71236 to replace dSpCas9-KRAB domain with ^VP64^dSpCas9^VP64^. This plasmid was further modified to generate an all-in-one CRISPRa lentiviral plasmid expressing ^VP64^dSpCas9^VP64^, a gRNA scaffold, and Thy1.1 by replacing PuroR with Thy1.1. The gRNA expression plasmid was generated by modifying Addgene plasmid #83925 to contain an optimized gRNA scaffold^32^ and a puromycin resistance cassette in place of Bsr. Individual gRNAs were ordered as oligonucleotides (IDT), phosphorylated, hybridized, and cloned into the plasmids using BsmBI sites. ORF overexpression isoform plasmids were constructed by inserting FOXO4 ORFs into a lentiviral plasmid encoding hUBC-ORF-2A-mCherry.

### Human astrocyte culture

Primary human astrocytes derived from the fetal cerebral cortex were purchased (Sciencell 1800) and routine cell culture was performed following the supplier’s protocol. Upon thawing, cells were seeded on plates coated with poly-D-lysine (PDL), and grown in Astrocyte Medium (Sciencell 1801) supplemented to 10% fetal bovine serum (Gibco) to encourage proliferation. Culture medium was replaced every two days until cultures reached 70% confluence, and subsequently every three days until cultures reached 95% confluence. To passage, cells were rinsed with PBS and dissociated using 0.025% trypsin. To eliminate residual potential progenitor cells and non-proliferative neuronal cells, astrocytes expanded for two passages. All experiments were initiated upon replating after the second passage reached confluence. Astrocytes were subjected to freeze/thaw cycles.

### Human astrocyte transduction and state reprogramming

Primary human astrocytes were cultured as described above until the second passage reached confluence. Cells were then dissociated, counted, and transduced by plating at 9 x 10□ cells/cm^2^ in fresh Astrocyte Medium containing packaged lentivirus encoding CRISPRa machinery, ORF, or shRNA. The day of transduction was designated as day 0. On day 1, medium was replaced with fresh astrocyte medium without lentivirus. Beginning on day 2, 1 µg/mL puromycin (Thermo Fisher) was added to all medium changes through day 6 to select for transduced cells. To support the survival of diverse neural cell states, culture medium was switched to a basal co-culture formulation on day 3 consisting of DMEM/F12 supplemented with 0.5% FBS, 1x N-2, 3.5 mM glucose, and 100 U/mL penicillin-streptomycin (all from Gibco). On day 8, the co-culture medium was supplemented with 20 ng/mL BDNF and 10 ng/mL NT-3 (Peprotech). Unless otherwise noted, all cell culture media was refreshed every 48 hours. Unless otherwise noted, cells were collected for analysis on day 10.

### Lentiviral production and titering

Lentivirus was packaged following the approach of McCutcheon et al.^33^ with minor modifications. Briefly, HEK293T producer cells were plated in OptiMEM Reduced Serum Medium (Gibco) supplemented with 1x GlutaMAX (Gibco), 5% fetal bovine serum (Gibco), 1 mM sodium pyruvate (Gibco), and 1x MEM non-essential amino acids (Gibco). Approximately 18 hours later, HEK293T cells were transfected with the second-generation lentiviral packaging and envelope plasmids pMD2.G and psPAX2 together with the transgene vector using Lipofectamine 3000 (Thermo Fisher). Culture medium was replaced 6 hours post-transfection, and viral supernatant was harvested and pooled at 24 and 48 hours post-transfection. The collected media containing the packaged lentivirus was centrifuged to remove cell debris and concentrated 50-fold with Lenti-X Concentrator (Takara). Lentiviral titers were measured as described by Gordon et al.^34^. Briefly, astrocytes were transduced with serial dilutions of virus, the medium was changed after 24 hours, and after four days cells were washed four times with PBS. Genomic DNA was then isolated using a DNeasy Blood & Tissue Kit (Qiagen), and integrated viral copies were quantified by qPCR with primers for genomic DNA (LP34), integrated viral DNA (WPRE), and unintegrated plasmid backbone (BB). Viral doses that led to cytotoxicity or cell morphology changes were excluded from analysis.

### Intracellular flow cytometry

To measure intracellular MAP2 expression by flow cytometry, astrocytes were washed with PBS, detached using 0.025% trypsin, dispersed into single-cell suspensions, and fixed with Intracellular Fixation Buffer (eBioscience) with gentle rocking at room temperature for 20 minutes. After fixation, cells were permeabilized with Intracellular Permeabilization Buffer (eBioscience). They were then washed 3x with Permeabilization Buffer and incubated for 10 minutes at room temperature in permeabilization buffer containing 0.2 M glycine (Sigma) and 2.5% FBS (Gibco) to block nonspecific binding. Blocked cells were then stained for 30 minutes at room temperature in the dark with gentle rocking using a rabbit anti-MAP2 antibody conjugated to Alexa Fluor 488 (Abcam, ab225316). Stained cells were rinsed three times with Permeabilization Buffer and once with FACS buffer (PBS with 0.5% BSA and 2mM EDTA), resuspended in FACS buffer, then analyzed and/or sorted on a Sony SH800z cell sorter.

### RT-qPCR

To assess RNA levels of transcripts of interest, total RNA was isolated using a Total RNA Purification Plus Kit or Total RNA Purification Plus Micro Kit (Norgen). cDNA was synthesized using SuperScript VILO (Thermo Fisher) from an equal mass of input RNA. cDNA was diluted before quantification. Quantitative PCR was performed with Perfecta SYBR Green Fastmix (Quanta BioSciences) on a CFX96 Real-Time PCR Detection System (Bio-Rad). All RT-qPCR primer sets were validated with standard curves and melt curves to assess efficiency and specificity. All RT-qPCR data is presented as log2 fold change normalized to GAPDH unless otherwise noted.

### Immunocytochemistry

To examine protein expression and morphology of transduced cells, transductions and all following procedures were conducted on PDL-coated 8-well µ-Slide (Ibidi Bioscience). At day 10, cells were rinsed gently with PBS twice and fixed with 4% formaldehyde (Pierce) for 15 minutes at room temperature. Fixed cells were then rinsed 3x with PBS, permeabilized with PBS supplemented with 0.1% Triton-X (Sigma), rinsed with PBS, and blocked with PBS supplemented with 0.1% Tween-20 (Sigma) and 10% Normal Goat Serum (Sigma). Incubation with primary antibodies was carried out at 4°C overnight. Primary antibodies employed in this study: Rabbit anti-MAP2 (1:500 dilution, Millipore Sigma ab183830), mouse anti-NeuN (1:1000 dilution, Millipore Sigma MAB377). After primary antibody incubation, cells were rinsed 3x with PBS supplemented with 0.1% Tween-20 (Sigma). Secondary staining: Cells were incubated with DAPI (Invitrogen) and cross-adsorbed secondary antibodies (Invitrogen) conjugated to Alexa Fluor 488 or 647 for 1 hour in the dark at room temperature. Stained cells were rinsed 3x with PBS and imaged with a Zeiss 780 upright fluorescent microscope. Images were taken at matched exposures and post-processing was conducted with the same parameters in parallel.

### RNA-sequencing

Total RNA was extracted as described above for RT-qPCR, and concentrations were determined prior to submission to Azenta for standard RNA-seq with poly(A) selection and ERCC spike-in controls. Libraries were sequenced on an Illumina platform using a 150-cycle paired-end kit. Raw FASTQ reads were trimmed with Trimmomatic v0.32^35^ and aligned to the GRCh38 reference genome using the STAR RNA-seq aligner v2.4.1a^36^. Gene-level counts were generated with featureCounts from the Subread package v1.4.6-p4^37^, using Gencode v22 for transcript annotation. Differential expression was assessed in DESeq2^38^, which applies a negative binomial generalized linear model and evaluates significance with the Wald test. Upregulated differentially expressed genes were then submitted to EnrichR^39^ for gene-ontology and functional annotation using the GO Biological Processes 2023 database.

### Design of gRNA libraries

Paired gRNA screen: Active gRNA hits with significant effects on astrocyte state were extracted from a previous screen of all transcription factor-encoding genes^40^. The resulting library consisted of 119 gRNAs targeting 90 TFs and 14 non-targeting gRNAs. FOXO4 promoter screen: SpCas9 gRNAs within 1kb of the FOXO4 TSS were generated with GuideScan2^41^. All gRNAs with a specificity score > 0.2 were retained for screening. The resulting library consisted of 199 targeting guides and 41 non-targeting gRNAs. Mismatch screen: Six FOXO4-targeting gRNAs with a range of on-target activation levels were selected for comprehensive screening of variants. In total, 168 variants for each gRNA were included for screening, including the original gRNA, all single base mismatches along the 20bp protospacer, 1- and 2-bp truncations from the PAM-distal end, 1bp truncation with all single base mismatches in the 5 PAM-distal bases, and all double mismatch combinations in the 5 PAM-distal bases. To this set, we added 100 non-targeting gRNAs and 6 MAP2-targeting positive controls. Then, variants with poly Ts and/or BsmBI sites were excluded, for a total of 1107 gRNAs. Secondary structure predictions for mismatched screen sgRNA variants were generated with ViennaRNA (v2.x) RNAfold at 37 °C^42^.

### Pooled CRISPRa screening

Pooled sgRNA libraries were cloned into the all-in-one CRISPRa lentiviral vector as described previously^33^, packaged into lentivirus and delivered to astrocytes as described above at MOI = 0.3 for all screens with the exception of the paired screen, in which the screened library was delivered at MOI = 0.3 but gRNA-hit was delivered at an MOI = 3, with Thy1.1 in place of the puromycin resistance cassette. After transduction, we carried out the human astrocyte state reprogramming media change paradigm described above. On day 10, cells were collected with 0.025% trypsin, fixed, permeabilized, stained, and sorted based on intracellular MAP2 expression as described above. To identify gRNAs that result in astrocyte state reprogramming, we sorted the lower and upper 10% bins of cells based on MAP2 expression for gRNA library amplification, construction and sequencing. All screen replicates were sorted at a minimum of 200x gRNA coverage (each gRNA was represented by at least 200 cells). After sorting, cells were lysed and DNA crosslinking was reversed by incubation at 65°C overnight with Proteinase K using a PicoPure DNA Extraction Kit (Arcturus). After reverse crosslinking, nucleic acids were purified by ethanol precipitation and dsDNA was quantified with a Qubit Fluorometer. Lentivirally-integrated gRNA cassettes from each cell population were then amplified from genomic DNA with barcoded custom i5 and i7 primers for Illumina sequencing. Amplicons were isolated from gDNA and primers with a double-sided SPRI bead selection, pooled, and sequenced on an Illumina MiSeq or NextSeq using custom Read 1 and index primers. To analyze differential gRNA abundance in MAP2 expression bins, FASTQ files were aligned to a gRNA library FASTA using Bowtie2^43^, and matrices containing gRNA counts in each MAP2 bin were analyzed using DESeq2^38^. gRNAs with DESeq2 Padj < 0.01 were designated as hits.

### Perturb-seq

Paired screen gRNA library was delivered at MOI = 0.3 alongside gRNA-hit (MOI = 3) with Thy1.1 in place of the puromycin resistance cassette. Ten days after transduction, cells were collected and gRNA and gene expression libraries were prepared using the 10X High-throughput kit with 5’ gRNA Direct Capture (10x Genomics) according to manufacturer protocol and sequenced on an Illumina Novaseq. Demultiplexing and unique molecular identifier (UMI) count generation for each transcript and gRNA per cell barcode was performed using CellRanger v6.0.1 (10x Genomics). UMI counts tables were extracted and used for subsequent analyses in R using Seurat v4.1.0^44^ and normalized with sctransform^45^. High-quality cells across donors were aggregated for further analyses. gRNA assignment: gRNAs were assigned to cells using CLEANSER^46^. Cells were then grouped for differential expression analysis using MAST^47^ based on gRNA assignments. A stringent threshold for gRNA-hit counts was applied to ensure all cells had received gRNA-hit, and all cells below the threshold were excluded from downstream analysis. DE testing: For differential gene expression analysis, for each gRNA, cells that received a given TF-targeting gRNA alongside gRNA-hit were compared to cells that received a non-targeting gRNA alongside gRNA-hit. DE testing was performed using Seurat’s FindMarkers function with the hurdle model implemented in MAST^47^. Upregulated DE genes were input into EnrichR’s GO Biological Process 2023 database for functional annotation as described above.

### ChIP-seq

ChIP experiments were performed in triplicate and harvested 8 days after transduction. For each replicate, 2 x 10^7^ astrocytes were resuspended in Astrocyte Medium with 1% formaldehyde (Pierce) and fixed for 10 minutes at room temperature with gentle rocking. Fixation was blocked with 0.125 M glycine for 5 minutes. Cells were then washed twice with ice-cold PBS and resuspended in cell lysis buffer (5 mM HEPES, 85mM KCl, 0.5% NP-40 pH 8.0) with protease inhibitor. To facilitate nuclei extraction, cells were passed 15 times through a Dounce homogenizer before incubation for 15 minutes at 4°C. Nuclei were then resuspended in nuclear lysis buffer (50 mM Tris HCl, 10 mM EDTA, 0.1% SDS pH 8.1) and sonicated to 200-700 bp fragments using a Diagenode Bioruptor Pico for 15 cycles of 30□seconds on and 30□seconds off. Before IP, a sample was collected for input control. For each replicate, we conjugated 5 µg monoclonal Anti-FLAG M2 (Sigma F1804) to 200 µl Dynabeads M-280 Sheep Anti-Mouse IgG (Invitrogen 11201D). Sheared chromatin was then incubated with antibody-conjugated beads on a rotator overnight at 4°C. Beads were washed five times with a LiCl wash buffer (100 mM Tris pH 7.5, 500 mM LiCl, 1% NP-40, 1% sodium deoxycholate), and remaining ions were removed with a TE wash (10 mM Tris-HCl, pH 7.5, 0.1 mM EDTA). Chromatin fragments were eluted in an IP elution buffer (PBS, 1% SDS, 0.1M NaHCO3) at 65°C for 1 hour. Supernatant containing sheared chromatin was incubated at 65°C overnight to reverse DNA-protein crosslinks. ChIP DNA was purified with QIAquick PCR Purification kit (Qiagen). ChIP-seq libraries were constructed with a KAPA HyperPrep kit (Roche). ChIP DNA was quantified and amplified with 7 PCR cycles after KAPA adapter ligation. Libraries were pooled and sequenced with 1% PhiX spike-in on an Illumina NextSeq with a P4 XLEAP-SBS reagent kit.

### ChIP-seq analysis

Reads that aligned to the integrated lentiviral cargo were removed using Bowtie2^43^. Remaining reads were then aligned to the hg38 reference genome, and PCR duplicates were removed using rmdup from SAMtools^48^. Peaks were called in each replicate individually using MACS2^49^. Peaks passing qval < .05 not in hg38 blacklist regions were then input to IDR analysis^50^ to identify high-confidence reproducible peaks across replicates. Peaks passing a stringent IDR threshold (IDR < 0.001) in all replicate comparisons within a condition were merged using BEDtools^51^ to generate per-gRNA peaksets, then a union peakset was generated by merging across gRNA conditions. A reads in peaks counts matrix was generated using featureCounts from Subread^37^. Peak counts from gRNA-hit and non-targeting samples were compared using DESeq2^38^ with non-variable genomic regions used as controlGenes for normalization. To generate the set of non-variable regions, we binned the hg38 genome into bins matching the average peak length, removed bins overlapping peaks or hg38 blacklist regions, and randomly sampled a set of 50,000 GC content-matched bins for read counting and subsequent normalization. Peaks were annotated with the nearest gene within 50kb for intersection with RNA-seq data using BEDtools closest^51^.

To generate browser tracks, BAM files were converted to normalized bigWig tracks using deepTools^52^. For each sample, reads were extended to estimated fragment length and binned at 25 bp resolution. Genome coverage was computed with bamCoverage, applying read-per-genome-coverage (RPGC) normalization to an effective hg38 genome size of 2.91 Gb and masking ENCODE hg38 blacklist regions. Tracks were then visualized using the Integrative Genomics Viewer^53^. Enriched motifs in peaksets were identified with HOMER^54^. Consensus peaks were analyzed using findMotifsGenome.pl against the hg38 reference genome. *De novo* motif enrichment was computed within 200 bp of each peak center. Consensus peaks were annotated with ChIPseeker using the UCSC hg38 knownGene transcript database. Peaks were assigned hierarchically to the most specific feature, prioritizing Promoter > 5′ UTR > 3′ UTR > Exon > Intron > Enhancer > Intergenic. A ±1 kb window around TSS defined promoters.

### *In vitro* dCas9 binding assay

DNA library designs: High-throughput DNA libraries were built to profile dCas9 binding specificity for 10 gRNAs (gRNA-hit, g1, g23, g110, g2, g25, g28, g113, g155, g194) targeting the promoter and 5′ UTR of FOXO4 (guide sequences in Table S2). The library comprised two main probe types: (i) WT PAM probes, representing on-target sites that include the 20nt protospacer and the 3nt PAM, each embedded in its native genomic context with an additional 8nt flanking the PAM and 5nt flanking the protospacer; and (ii) No PAM controls, in which the PAM in the corresponding WT PAM sequence was changed to TAC to assess nonspecific binding. For each WT PAM target, every possible single nucleotide substitution across the target region was introduced individually. For gRNA-hit, the design also incorporated all IDT-predicted NGG off-target sites containing ≤3 mismatches.

DNA array preparation and RNP:DNA binding assay: Libraries were synthesized on Agilent microarray slides in 16×25k or 24×13k formats (i.e., 16 or 24 chambers with 25,000 or 13,000 spots per chamber, respectively). Each unique sequence was printed in eight replicates randomly distributed within a chamber. Oligos were 60nt single-stranded DNAs consisting of a 36nt variable region followed by a 24nt constant tail (5′-GTCTGTGTTCCGTTGTCCGTGCTG-3′) complementary to a primer used to convert features to double-stranded DNA by primer extension, as described previously. Arrays were incubated in an extension mix containing 26 mM Tris-HCl (pH 9.5), 6.5 mM MgCl□, 1.16 µM primer, 163 µM dNTPs, and 32 units Thermo Sequenase DNA polymerase, using the following temperature program: 85 °C for 10 min, 75 °C for 10 min, 65 °C for 10 min, and 60 °C for 90 min.

Ribonucleoprotein (RNP) complexes were assembled by incubating 2 µM dSpCas9-His (IDT Alt-R™ S.p. dCas9 Protein V3) with 4.8 µM sgRNA (IDT Alt-R™ CRISPR Custom Guide RNAs) for 10 min at 37 °C in 20 mM Tris-HCl (pH 7.4), 150 mM KCl, 10% glycerol, 5 mM MgCl□, and 1 mM TCEP. Following a universal protein-binding microarray (PBM) workflow, double-stranded arrays were blocked with 2% (w/v) nonfat milk (Sigma) for 30 min and then incubated for 1 h at room temperature with 100 nM RNP in binding buffer (20 mM HEPES-KOH, pH 7.5, 150 mM KCl, 5 mM MgCl□, 5% glycerol, 0.025% Triton X-100, 1 mM TCEP). After binding, arrays were stained with 10 µg/mL Penta·His Alexa Fluor 488 Conjugate (Qiagen) to detect bound RNPs. Fluorescence was quantified on a GenePix 4400A microarray scanner using GenePix Pro 7.0 software. For each sequence, the median fluorescence intensity across its eight replicate spots was used for analysis.

### FOXO4 shRNA and ORF overexpression

shRNA expression: shRNA targeting FOXO4^55^ was cloned into the lentiviral gRNA expression plasmid in place of the gRNA cassette and delivered to astrocytes alone or alongside gRNA-hit cloned into the all-in-one CRISPRa lentiviral plasmid. ORF overexpression: ORFs encoding the 1515bp canonical codon-optimized FOXO4 ORF isoform (ENST00000374259) was ordered from IDT. Non-codon-optimized protein-coding ORF isoforms (ENST00000374259, ENST00000341558) were amplified from the Multiplexed Overexpression of Regulatory Factors (MORF) Library^56^ (Addgene #192821) and delivered to astrocytes alone or alongside gRNA-hit cloned into the all-in-one CRISPRa lentiviral plasmid. After transduction, we carried out the human astrocyte state reprogramming media change paradigm described above and lysed cells for RNA purification at day 10.

## RESULTS

### Robust reprogramming of astrocyte cell state with CRISPRa

We explored the dynamics of CRISPRa gRNA on- and off-target activity in the context of astrocyte cell state reprogramming. Astrocytes are multifunctional glial cells that undergo complex state transitions in response to injury and neurodegeneration^57^. As they have latent proliferative potential throughout mammalian adulthood, astrocytes have also gained significant attention as starting material for cell reprogramming-based neuroregenerative therapies, such as astrocyte-to-neuron reprogramming^58–67^. Therefore, they represent a cell type amenable to functional genomic CRISPR screening and a potential target for CRISPRa/i-based therapies. As such, astrocyte transcriptional states and the factors that regulate them, whether natural states associated with reactive gliosis or engineered states designed for neuroregenerative approaches, deserve further study.

Previously we conducted a high-throughput CRISPRa screen of all genes encoding human TFs (TFome) to identify factors that could reprogram astrocytes to neuron-like states^40^. This screened employed MAP2, a specific pan-neuronal marker gene, as a proxy for cells driven from astrocyte state to a neuron-like transcriptional state^68^, and identified a single gRNA designed to target the promoter of FOXO4 (referred to as “gRNA-hit”) as a potent astrocyte state reprogramming factor. However, despite the strong effect size of gRNA-hit, the other 5 gRNAs targeting FOXO4 did not emerge as hits^40^. Additionally, FOXO4 has not previously been associated with astrocyte homeostasis or astrocyte state reprogramming. While FOXO4 is necessary for differentiation of human embryonic stem (ES) cells to a neural lineage^69^, it is more commonly associated with cell senescence and aging^70^. Therefore, we first profiled the effect of FOXO4 activation by gRNA-hit in astrocytes to gain insight into gRNA-hit-driven CRISPRa activity and transcriptional state reprogramming.

To test whether CRISPR activation with gRNA-hit indeed has significant effects on astrocyte gene expression, we transduced primary human astrocytes from 3 donors with lentivirus encoding ^VP64^dSpCas9^VP64^ and gRNA-hit and measured the expression of neural lineage marker genes after 10 days. We first measured the expression of MAP2 and observed conversion of the astrocyte population to >50% MAP2-positive cells (**Figure 1A, Figure S1A, S1B**). We also observed a ∼360-fold upregulation of NeuN (encoded by *RBFOX3*), a nuclear marker of post-mitotic neurons classically used to identify neurons *in vivo* (**Figure 1B, C**), supporting that gRNA-hit induced an astrocyte state change.

**Figure 1:**
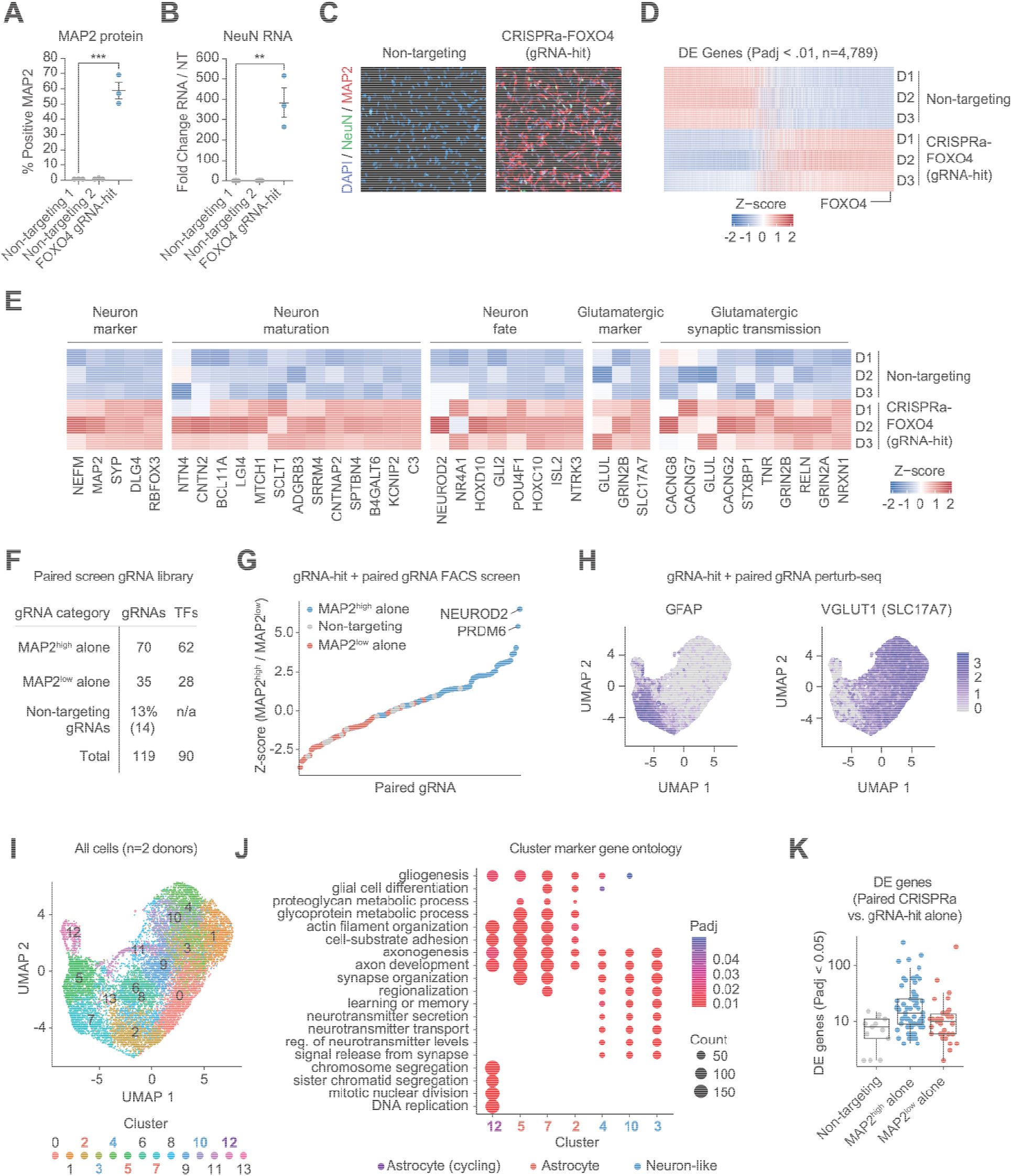
Robust reprogramming of astrocyte cell state with CRISPRa. A. MAP2 protein level after CRISPRa-FOXO4 with gRNA-hit, as proxy for astrocyte cell state change. B. NeuN RNA (RT-qPCR) level after CRISPRa-FOXO4 with gRNA-hit. In A and B, **p < 0.01, ***p < 0.001 by global one-way ANOVA with Dunnett’s post hoc test comparing all groups to non-targeting 1; error bars represent SEM. C. MAP2 and NeuN immunofluorescence staining after CRISPRa-FOXO4 with gRNA-hit, 10 days post-transduction. D. Differential expression (DE) analysis of RNA sequencing of ^VP64^dSpCas9^VP64^ + gRNA-hit vs. ^VP64^dSpCas9^VP64^ + non-targeting gRNA in astrocytes from 3 donors, 10 days post-transduction. FOXO4 is labeled. DE genes are determined using a Wald test by DESeq2 (Padj <0.01). E. Z-score of key genes of interest corresponding to gene ontology terms categories. DE genes are determined using a Wald test by DESeq2 (Padj <0.01). F. Summary of paired gRNA library. G. Z-score of screened gRNAs in paired gRNA screen, ranked from low (enriched in MAP2^low^ cell population) to high (enriched in MAP2^high^ cell population). H. Feature plots displaying count distribution of astrocyte (left) and glutamatergic neuron (right) marker genes in perturb-seq UMAP embedding. I. UMAP embedding from paired gRNA-hit perturb-seq. J. Selected enriched biological processes for clusters of interest. Statistical significance was determined using a two-tailed Fisher’s exact test followed by Benjamini–Hochberg correction. K. Numbers of DE genes associated with each perturbation (significant gRNA-gene links), as defined using a two-tailed MAST test with Bonferroni correction.

To more widely assess the transcriptomic effect of CRISPRa with gRNA-hit, we delivered the same CRISPRa construct to astrocytes from 3 donors and performed RNA-seq. We expected that modulating the expression of a TF such as FOXO4, which in turn regulates the expression of other genes, would result in many genes being differentially expressed (DE). Indeed, CRISPRa with gRNA-hit had a striking effect on astrocyte gene expression, with 4,789 genes differentially expressed (FDR < 0.01) (**Figure 1D**). Among DE genes, FOXO4 was significantly upregulated, as expected. We found that the set of upregulated genes also included canonical neuron marker genes, as well as genes associated with neuron maturation and acquisition of neuronal fate (**Figure 1E**). Gene sets associated with a glutamatergic neuronal subtype were also significantly upregulated, including canonical glutamatergic marker genes *SLC17A7* (VGLUT1), *SLC17A6* (VGLUT2), and *GRIN2B* (a component of the ionotropic glutamate receptor), as well as genes involved in glutamatergic synaptic transmission^71^. Together, these data suggest that CRISPRa with ^VP64^dSpCas9^VP64^ and gRNA-hit potently reprograms astrocyte transcriptional state and upregulates gene sets associated with excitatory neurons.

We next conducted paired CRISPRa screens to test whether additional TFs were capable of enhancing gRNA-hit’s astrocyte state reprogramming. We screened a gRNA library consisting of all gRNA hits from our previous TFome screen^40^, including gRNAs enriched in the MAP2^high^ cell population (pro-neuronal hits) and gRNAs enriched in the MAP2^low^ population as controls (**Figure 1F**). We transduced astrocytes with lentivirus encoding ^VP64^dSpCas9^VP64^ and gRNA-hit at MOI = 3, and concurrently delivered a lentivirus encoding our gRNA library at MOI = 0.3 such that after antibiotic selection, almost all cells would have received ^VP64^dSpCas9^VP64^, gRNA-hit, and one other gRNA. We first sorted cells based on MAP2 expression via FACS. Overall, TFs that decreased MAP2 expression alone also decreased MAP2 expression when paired with gRNA-hit, and TFs that increased MAP2 expression alone also increased MAP2 expression when paired with gRNA-hit (**Figure 1G**). We next performed a Perturb-seq screen^72^, conducted identically to the FACS-based screen, but rather than sorting cells based on MAP2 expression, we performed scRNA-seq. In total, we profiled ∼16,000 high-quality cells from 2 donors (**Figure S1C, D**). The majority of profiled cells expressed genes associated with mature glutamatergic neurons, further supporting significant reprogramming of astrocyte state (**Figure 1H, Figure S1E**). In total, unsupervised clustering resulted in 14 cell clusters (**Figure 1I**), with gene ontology analysis of cluster marker genes supporting the presence of distinct transcriptional states (**Figure 1J**). We next examined the effect of each paired perturbation alongside gRNA-hit by performing differential expression analysis comparing the transcriptomes of cells that received gRNA-hit with a non-targeting gRNA to transcriptomes of cells that received gRNA-hit with each targeting gRNA individually. The effects of pairing targeting gRNAs with gRNA-hit were modest compared to gRNA-hit alone (mean DE genes = 28, 18, and 8 for positive hits, negative hits, and non-targeting gRNAs respectively) (**Figure 1K**). Nevertheless, identified DE genes were consistent with previous TF associations. For example, genes upregulated by activation of NEUROD2 were largely related to neuronal processes (**Figure S1F**). Overall, CRISPRa with gRNA-hit alone and in combination with other TF-targeting gRNAs has a striking impact on the astrocyte transcriptome, reprogramming cells into a distinct state with gene sets associated with glutamatergic neurons. While this phenotype can be modestly modulated by paired activation of other TFs, the final transcriptional state appears dominated by the effects of CRISPRa with gRNA-hit.

### gRNA-hit has unique effects among SpCas9 gRNAs targeting the FOXO4 promoter

CRISPRa with gRNA-hit has a potent effect on astrocyte cell state, changing expression of thousands of genes. However, gRNA-hit was the only gRNA designed to target FOXO4 that emerged as a hit in the TFome screen; the 5 other screened gRNAs from the same optimized library^73^ were not enriched in the MAP2^high^ population^40^. Efficacy and efficiency of CRISPRa gRNAs depend on a myriad of factors including expression level, gRNA secondary structure, target site position relative to endogenous factors and regulatory elements, and chromatin context at the target site. Thus, we first hypothesized that the lack of screen hits from other FOXO4-targeting gRNAs was due to ineffective FOXO4 activation in astrocytes. To test this hypothesis, we individually expressed other screened FOXO4-targeting gRNAs in astrocytes via the lentiviral CRISPRa construct. However, all tested gRNAs activated FOXO4, from a range of 10-30-fold compared to non-targeting control (**Figure 2A**). Notably, only gRNA-hit robustly reprogrammed astrocyte state (as measured by MAP2 expression) despite gRNA-hit resulting in the *lowest* level of FOXO4 activation (∼8-fold) among tested FOXO4-targeting gRNAs.

**Figure 2:**
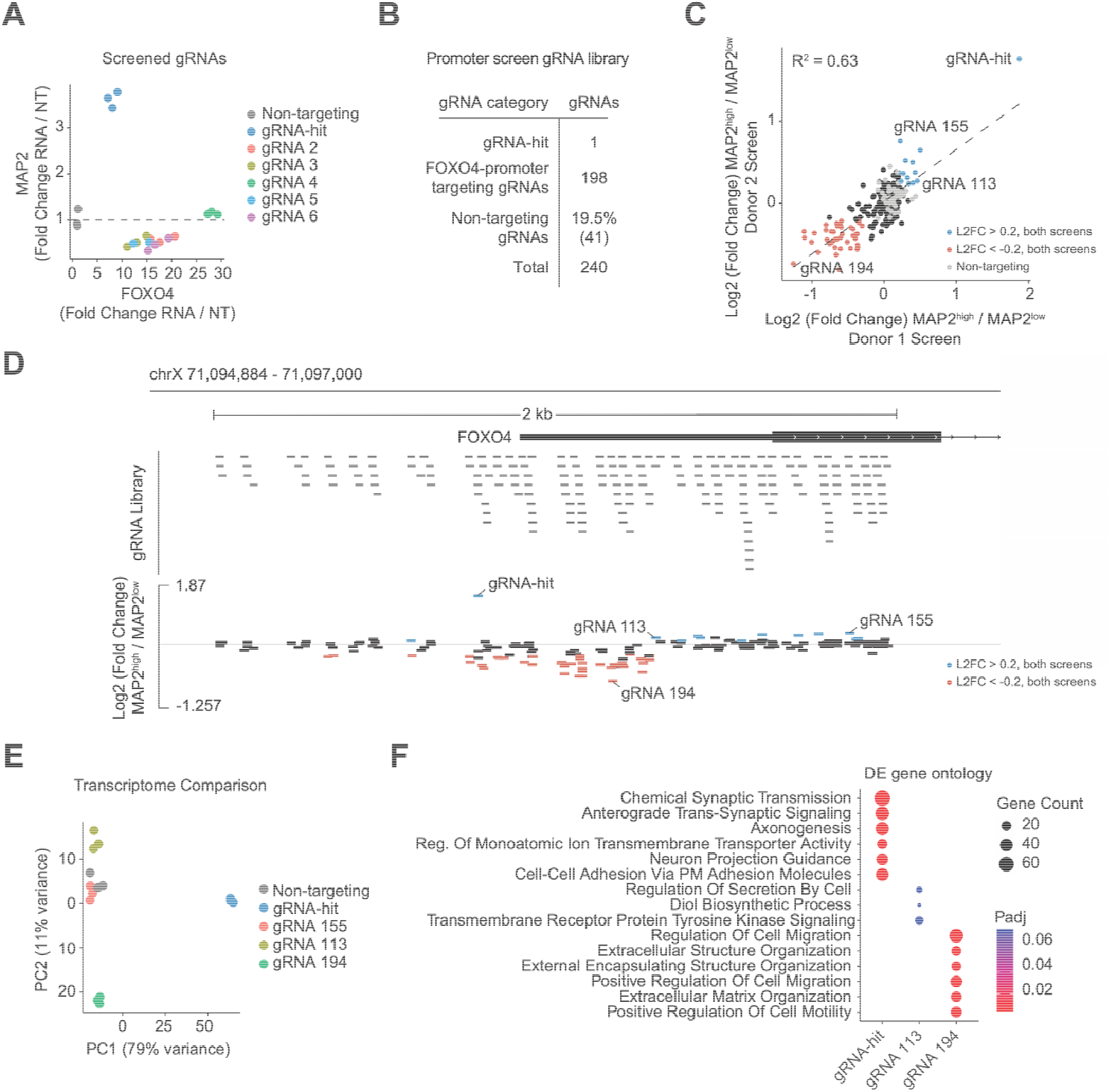
gRNA-hit has unique effects among FOXO4-targeting SpCas9 gRNAs. A. RNA levels of FOXO4 and MAP2 after delivery of 6 FOXO4-targeting gRNAs. B. Summary of FOXO4 promoter tiling gRNA library. C. Comparison of log2 fold change (MAP2^high^ / MAP2^low^) for screens conducted using astrocytes from 2 donors. D. FOXO4 promoter tiling screen library and fold changes from Donor 1 screen overlaid on FOXO4 promoter. E. Principal component analysis comparing resulting transcriptomes after delivery of gRNA-hit, gRNA 155, gRNA 113, gRNA 194, or non-targeting gRNA to primary astrocytes. F. Top enriched biological processes for upregulated DE genes after delivery of FOXO4-targeting gRNAs. No significant GO terms were associated with DE genes for gRNA 155. Statistical significance of term enrichment was determined using a two-tailed Fisher’s exact test followed by Benjamini–Hochberg correction.

To better understand the discrepancy between gRNA-hit and other tested FOXO4-targeting gRNAs and to attempt to identify other FOXO4-targeting gRNAs with robust reprogramming activity, we next tested every SpCas9 gRNA within 1kb of the FOXO4 transcriptional start site (TSS) via a FACS-based CRISPRa screen. We restricted the screened library to gRNAs with a Guidescan specificity score >0.2^74^. The screened library consisted of gRNA-hit, 198 other FOXO4-targeting gRNAs, and 41 non-targeting controls, for a total library size of 240 gRNAs (**Figure 2B**), which was screened in astrocytes from 2 donors. The screen revealed that while some gRNAs displayed modest positive effects on MAP2 (including gRNA 113 and 155) and others displayed modest negative effects on MAP2 (including gRNA 194), none rivaled the potency of gRNA-hit (**Figure 2C**). We did not discern any obvious dependence of position relative to the TSS on gRNA activity (**Figure 2D**).

CRISPRa/i gRNAs often modulate their target gene to different extents. We therefore reasoned that activation of the same factor (in this case FOXO4) by different gRNAs would lead to correlated effects (i.e. a similar set of downstream genes, but potentially with different fold changes). To examine if multiple gRNAs were driving the same general directionality of state change, we performed RNA-seq after delivery of positive hits (gRNA 113 & 155) or a negative hit (gRNA 194) that emerged from the FOXO4 promoter screen. Despite all targeting the promoter of FOXO4, these gRNAs displayed divergent effects with variable potency (**Figure 2E**), each yielding distinct DE gene sets (**Figure S2A**). Each perturbation successfully upregulated FOXO4 (**Figure S2B**).

However, gene ontology of other DE genes yielded pro-neuronal state terms for gRNA-hit, but not for other gRNAs (**Figure 2F**), and directionality of shared DE genes showed no clear correlation between gRNA-hit and other gRNAs (**Figure S2C**). Therefore, these data suggest gRNA-hit has unique effects compared to other FOXO4-targeting gRNAs.

### Mapping and characterizing genome-wide off-target gRNA activity

Given the unique effect of gRNA-hit among FOXO4 promoter-targeting gRNAs, we next profiled potential off-target activity of gRNA-hit to examine the possibility that its astrocyte state reprogramming could be driven by off-target effects. The Guidescan score^74^, for gRNA-hit fell in the top 40% of gRNA specificity scores for gRNAs in the subset of the Calabrese library that targets the TFome^73^, as well as our promoter tiling screen library (**Figure S3A, B**). However, CRISPR/Cas9 can display significant DNA and RNA off-target activity depending on Cas protein, editing modality, delivery modality, expression level and duration, and specific gRNA^75^. Thus, we performed ChIP-seq to rigorously assess on- and off-target binding of ^VP64^dSpCas9^VP64^ to the genome when complexed with gRNA-hit, and in parallel measured DNA-binding by dCas9 complexed with gRNAs of interest in vitro with a protein-DNA binding microarray assay^76,77^.

To map genomic binding of dCas9, we transduced primary astrocytes with lentivirus encoding our CRISPRa construct and either gRNA-hit or a non-targeting gRNA. Eight days after transduction, we immunoprecipitated chromatin bound to FLAG-tagged ^VP64^dSpCas9^VP64^ with an anti-FLAG antibody and sequenced bound DNA fragments (**Figure S3C**). As expected, we observed a strong peak in the promoter of FOXO4 at the intended gRNA-hit on-target site (**Figure 3A**). However, CRISPRa with gRNA-hit also displayed 14,386 other highly reproducible (IDR < 0.001) peaks across all three replicates (**Figure 3B**). These binding sites spanned genomic contexts. The majority of peaks (8,258 peaks, 57.3% of total) were in accessible chromatin, but CRISPRa with gRNA-hit also demonstrated robust binding to closed chromatin (6,129 peaks, 42.7% of total). Binding sites also spanned genomic elements: 2,744 (19.1%) were in gene promoters, while 6,046 (42.0%) were in intronic regions. Across these genomic elements, binding was stronger in accessible chromatin^78^ (**Figure 3C**). Further, though binding to the genome does not solely indicate effector activity, many of the bound regions proximal to genes were associated with upregulation of those genes (**Figure 3D**).

**Figure 3:**
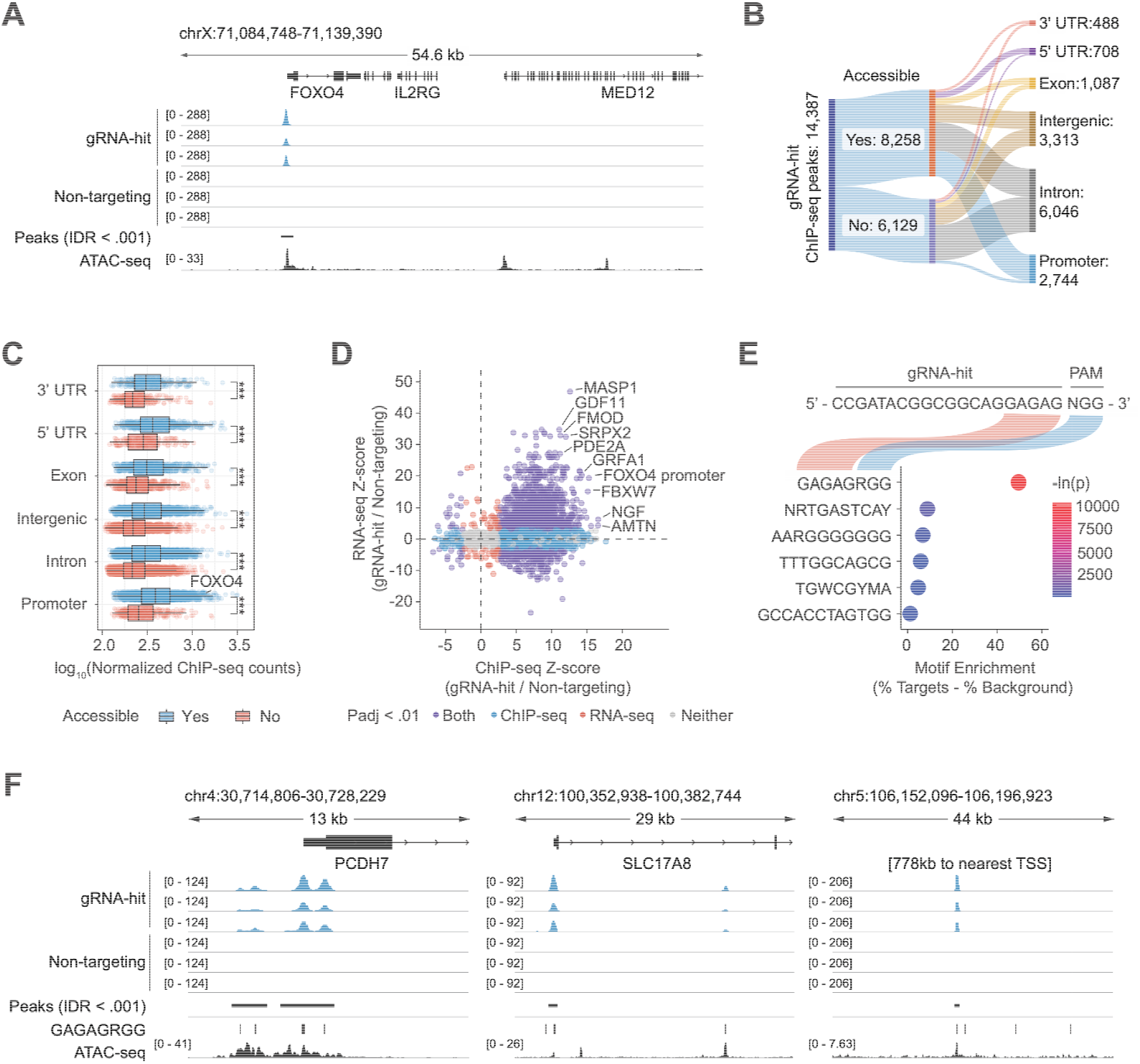
Mapping and characterizing genome-wide off-target gRNA activity. A. ChIP-seq peak demonstrating robust ^VP64^dSpCas9^VP64^ binding at the on-target site on the FOXO4 promoter after delivery of gRNA-hit. B. Sankey diagram of accessibility and genomic element type of 14,387 genomic binding sites. C. Normalized ChIP-seq counts by accessibility and genomic element type. Statistical significance was determined by Wilcoxon rank-sum test comparing accessible vs. non-accessible peaks within each feature followed by Benjamini–Hochberg correction, ***p < 0.001. D. Comparison of ChIP-seq peak DESeq2 Z-score with RNA-seq DESeq2 Z-score of proximal genes. Peak-gene links were defined as the closest gene within 50kb. E. HOMER de novo motif enrichment analysis in gRNA-hit ChIP-seq peaks. HOMER computes P values from the cumulative hypergeometric distribution and does not adjust for multiple hypotheses. F. Representative off-target gRNA-hit ChIP-seq peaks.

*De novo* DNA motif discovery in the gRNA-hit ChIP-seq peaks revealed GAGAGRGG as the most enriched motif, occurring in 56.7% of ChIP-seq peaks. This motif corresponds to the 5 PAM-proximal bases of gRNA-hit, followed by an SpCas9 PAM sequence (**Figure 3E**). While previous reports demonstrated the critical role of the 8-10bp gRNA seed region in DNA cleavage by Cas9 nuclease^31,79–82^, this finding supports that shorter PAM-proximal motifs can determine dCas9 binding^27,30^. However, not all instances of this motif in the genome were bound by dCas9, and not all ChIP-seq peaks contained the GAGAG motif followed by a matching PAM (43.3% of ChIP peaks did not contain GAGAGRGG) (**Figure 3F**, representative off-target sites). Thus, we concluded that gRNA-hit has pervasive off-target binding that results in many gene expression effects. This is in stark contrast to our previous dCas9 ChIP-seq analyses of highly specific gRNAs that recovered a single or very few genome-wide binding sites^25,29^.

### High-throughput mapping of gRNA substrate mismatch tolerance in vitro

Individual gRNAs have vastly different off-target propensities, but the determinants of gRNA binding specificity have not been fully defined^25,30^. Therefore, in parallel we sought to investigate the intrinsic properties of gRNA-hit that result in promiscuous binding. We used a protein-DNA binding microarray technology^83^ to measure DNA-binding by dCas9 complexed with gRNAs (**Figure 4A**). In this assay, dCas9 protein is first incubated with a gRNA to produce a ribonucleoprotein (RNP) complex, which is then incubated on a chip printed with a DNA library containing tens of thousands of custom-designed DNA sequences. To measure the DNA-binding specificity of the dCas9:gRNA-hit RNP, we designed a DNA library that included the gRNA-hit on-target site in its genomic sequence context, all possible single base-pair variants of the on-target site, a control sequence containing the wild-type target site but with the PAM mutated to a non-NGG control sequence (specifically, TAC), as well as putative genomic off-target sequences. Binding of dCas9 RNPs to these targets was measured with a fluorometric readout to assess sequence-specific and gRNA-dependent dCas9 binding specificity.

**Figure 4:**
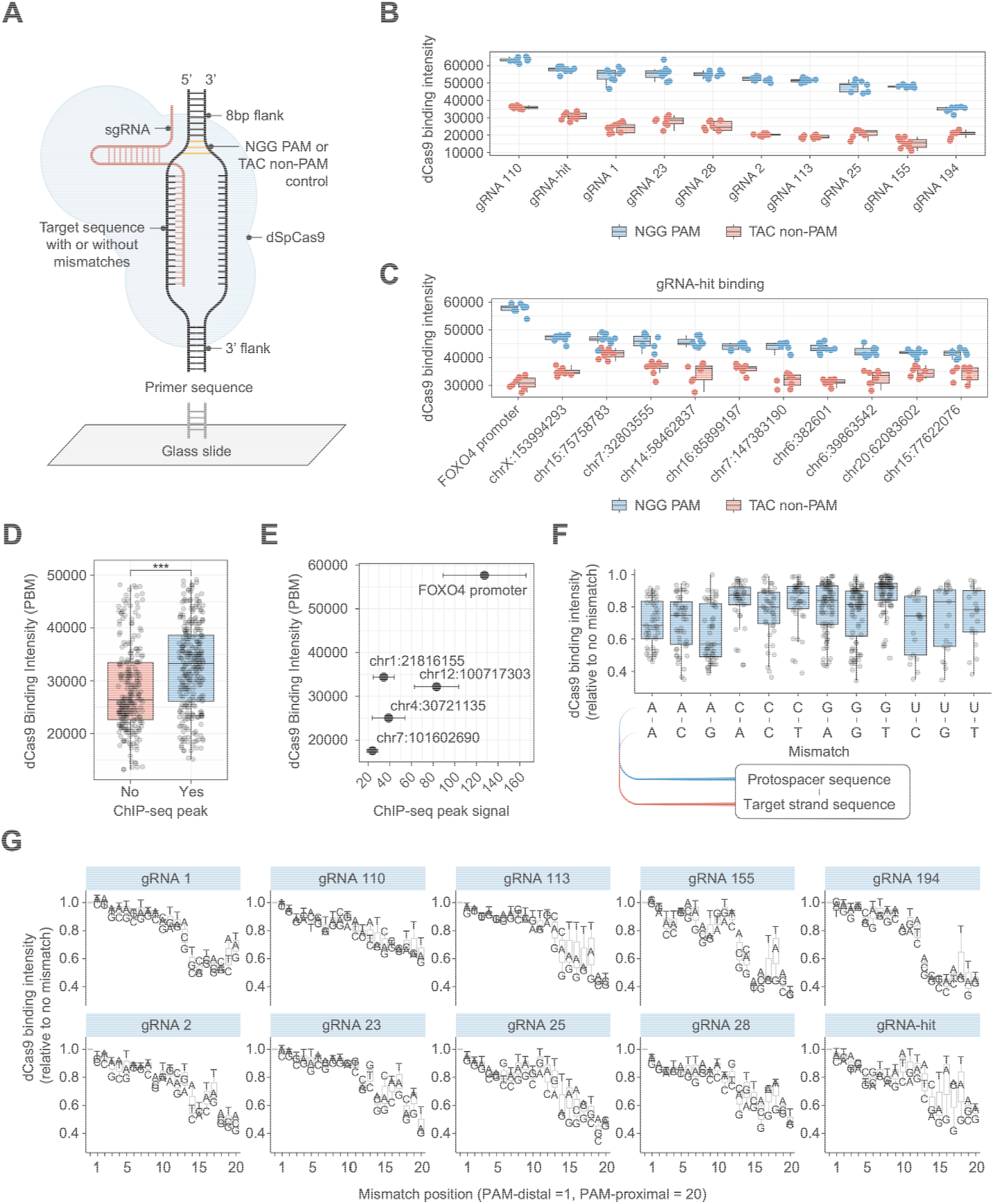
High-throughput mapping of gRNA substrate mismatch tolerance in vitro. A. Schematic of *in vitro* dCas9 binding assay. B. Binding intensities of 10 FOXO4-targeting gRNAs with on-target sequence adjacent to NGG PAM or TAC PAM. C. Binding intensity of dCas9 with gRNA-hit to on-target site and top 10 off-target sites of 66 tested, with either NGG PAM or TAC PAM. D. Binding intensity of dCas9 with gRNA-hit comparing binding intensity to sequences not in a ChIP-seq peak vs. sequences in a ChIP-seq peak. ***p < 0.001 by two-sided Wilcoxon rank-sum test. E. Scatterplot comparing *in vitro* binding intensities and ChIP-seq peak signal for peaks with MACS2 signalValue > 20. F. Protospacer-DNA-target mismatch tolerance. Binding is relative to gRNA-matched on-target binding. G. Protospacer-DNA-target mismatch tolerance by gRNA. Mismatch position = 1 corresponds to PAM-distal, mismatch position = 20 corresponds to PAM-proximal.

First, to validate the system, we designed DNA sub-libraries for 10 gRNAs from the FOXO4 promoter screen, including gRNA-hit, and compared binding of each gRNA to their wild-type on-target sites, with and without the PAM. All 10 gRNAs displayed robust binding to their on-target sequence containing the native PAM, relative to the on-target sequence containing a non-PAM ‘TAC’ sequence (**Figure 4B**). Next, we tested the binding affinity of dCas9:gRNA-hit to the on-target sequence in the FOXO4 promoter and 66 potential off-target sites in the human genome from *in silico* predictions based on sequence similarity. While dCas9:gRNA-hit displayed the strongest binding at the on-target site in the FOXO4 promoter, significant PAM-dependent binding was also observed at nominated off-target sites, with the highest affinity off-target site retaining ∼80% binding activity compared to the on-target sequence (**Figure 4C**). Comparison of the *in vitro* DNA array binding data to the ChIP-seq binding data revealed that gRNA-hit directed higher *in vitro* binding affinity to sequences that fell within off-target ChIP-seq binding sites (called as a peak by MACS2^49^) compared to other tested potential off-target sites not associated with a ChIP-seq peak, despite there being no difference in length or GC content (**Figure 4D, S4A**). Further, for sites with strong ChIP-seq signal, there was a generally positive correlation with *in vitro* binding affinity, demonstrating the fidelity of *in vitro* binding assays for modeling dCas9 off-target binding in cells (**Figure 4E**).

Next, we took advantage of the modularity and throughput of the DNA array platform to test gRNA protospacer-DNA target mismatches. We tested every single-base substitution across the DNA-target of 10 FOXO4-targeting gRNAs. DNA target strand mismatches containing a T in place of a native C, resulting in rG-dT mismatch, were more well-tolerated than other single-base mismatches, likely due to the ability to form a wobble base pair^84^ and consistent with previous findings^85–88^. We found that rA-dG mismatches were the least tolerated (**Figure 4F**). Upon segmentation of the 20bp protospacer into PAM-distal (10bp farthest from PAM) and PAM-proximal (10bp closest to PAM) regions, we observed that while rG-dT mismatches were well-tolerated throughout the protospacer, the effect size was larger in PAM-proximal bases, which displayed less mismatch tolerance overall (**Figure S4B**). Finally, for each gRNA, we examined mismatch tolerance at each base position individually (**Figure 4G, Figure S4C, S4D**). Across all gRNAs, PAM-distal mismatches were better tolerated than PAM-proximal mismatches. However, among all 10 tested gRNAs, gRNA-hit displayed the highest absolute tolerance level among PAM-proximal mismatches: dCas9 binding to substrates with a rG-dT wobble pair mismatch in position 16 or 18 of the gRNA-hit target strand displayed >90% binding activity compared to perfect substrate match. Additionally, gRNA-hit was among the most tolerant of mismatches in positions 10-13. Taken together, our *in vitro* binding assay results revealed that gRNA-hit has increased mismatch tolerance in critical PAM-proximal bases compared to other FOXO4-targeting gRNAs. This mismatch tolerance within the gRNA seed region is likely a major contributor to the multi-site, off-target binding across the genome which underlies gRNA-hit’s reprogramming capabilities.

### High-throughput mapping of promiscuous gRNA variants in cells

While PAM-proximal bases are critical for genomic binding, the specific sequence features that confer gRNA promiscuity remain unclear. To shed light on the relationship between gRNA sequence and promiscuity, we systematically tested mutated gRNA variants (mismatches in the gRNA protospacer rather than DNA target) in primary astrocytes. CRISPRa with gRNA-hit has widespread off-target activity (**Figure 3**) and a potent effect on MAP2 expression (**Figure 1A, E, S1A**,**S1B**), but other FOXO4-targeting gRNAs did not significantly impact MAP2 expression (**Figure 2 A,C,D**). We therefore once again employed MAP2 expression as a readout, this time as a measure of the transcriptional reprogramming driven by off-target activity of mutated variants of gRNA-hit (**Figure 5A**). We also comprehensively tested variants of 5 other FOXO4-targeting gRNAs as controls. We designed a gRNA library which included 168 variants for each protospacer. These variants include non-mutated protospacers (1 variant), all single base substitutions across the protospacer (60 variants), 1bp- and 2bp-truncations from the 5’ PAM-distal bases (2 variants), and a 1bp truncation alongside PAM-distal (5’) mismatches (15 variants). Previous work screening protospacer mismatches demonstrated that single mismatches are often tolerated and can be introduced to modulate dCas9 binding and CRISPRi activity. However, double mismatches are not well-tolerated in PAM-proximal bases^89^. Therefore, we also included all combinations of double-mismatches in 5 PAM-distal protospacer bases (90 variants). With non-targeting controls and MAP2-targeting positive controls, the library consisted of a total of 1,097 gRNAs.

**Figure 5:**
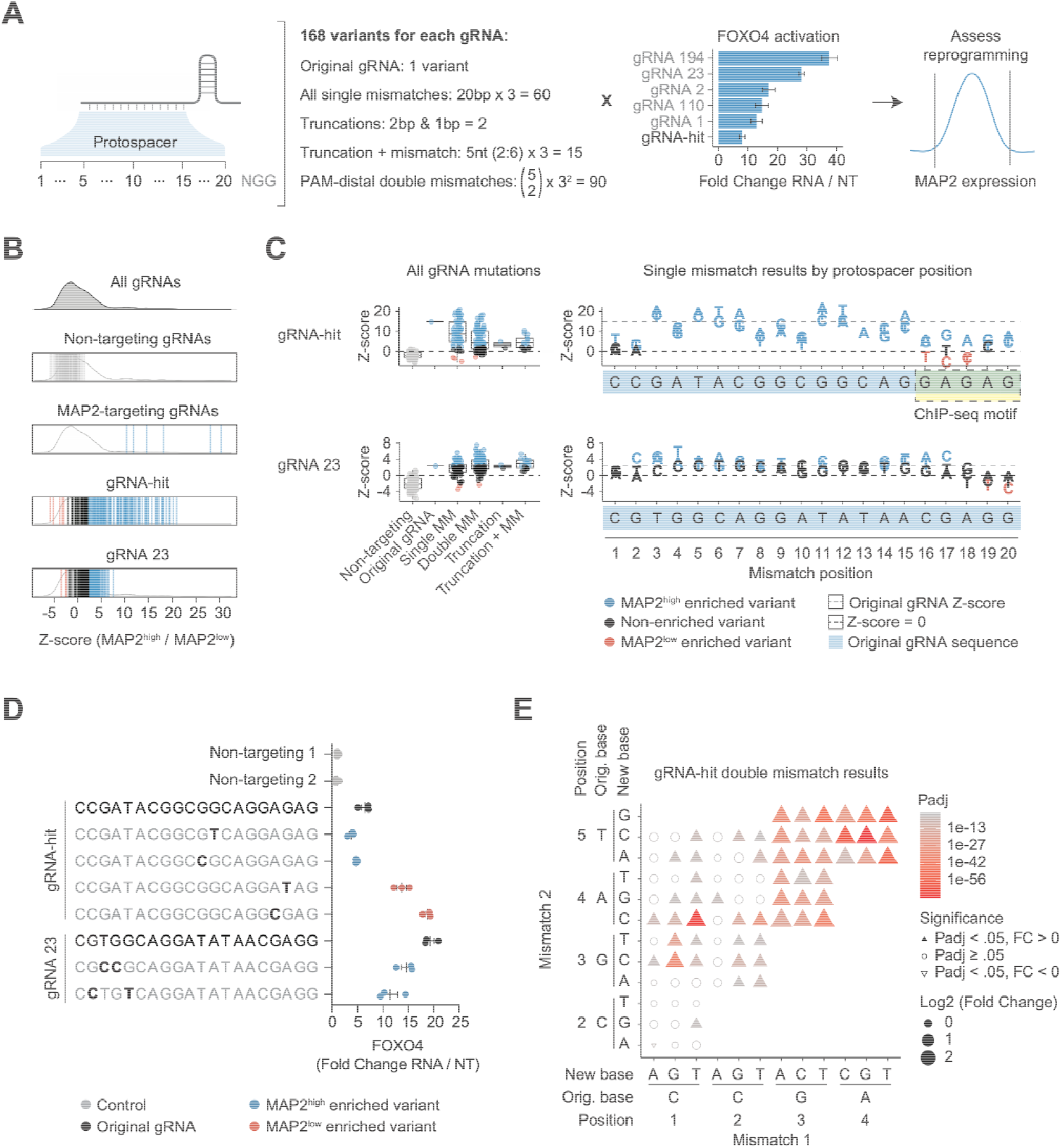
High-throughput mapping of promiscuous gRNA variants in cells. A. Schematic of protospacer mismatch screening approach. Six FOXO4-targeting gRNAs spanning a range of on-target activation levels were screened, with 168 variants per gRNA. A MAP2 FACS readout was employed to measure cell state reprogramming. B. Screen DESeq2 Z-score of gRNA variants. Enriched variants (Padj < 0.01) were defined using a paired two-tailed DESeq2 test with Benjamini–Hochberg correction. The overall distribution is slightly left-shifted due to strong positive fold change of gRNA-hit variants. C. gRNA variant screen DESeq2 Z-score by category (left) and single mismatch variants by base position (right) for gRNA-hit and control gRNA 23. Non-mutated gRNA Z-score is represented by a gray dashed line, Z-score = 0 is represented by a black dashed line, and plotted bases represent new DNA bases encoding the gRNA at each position. Non-mutated gRNA sequence is shown highlighted in light blue. All Z-scores in panels B and C refer to gRNA abundance in MAP2^high^ / MAP2^low^ cell populations. D. On-target FOXO4 activation levels of screened gRNA variants of interest. In B-D, non-targeting gRNAs are shown in gray, significant (Padj < .01) positive fold change gRNAs (enriched in MAP2^high^ cell population) are shown in blue, significant (Padj < .01) negative fold change gRNAs (enriched in MAP2^low^ cell population) are shown in red, and all other variants are shown in black. E. Screen significance and fold change for PAM-distal double mismatch variants of gRNA-hit, plotted by position of mismatches.

Results of this screen indicated that many gRNA-hit variants retained the ability to reprogram astrocyte state, implying the retention of off-target activity. (**Figure 5B**). Similarly, gRNA-hit reprogramming activity was retained after introduction of 5’ double mismatches and some truncated variants (F**igure 5C**, **left**). Mismatches introduced in positions 11 and 12 were particularly potent, consistent with our *in vitro* DNA array experiments which revealed strong mismatch tolerance at those positions (**Figure 5C, right**). However, introducing mismatches into other FOXO4-targeting control gRNAs only modestly modulated their effect on MAP2 expression (**Figure 5B**), or in other cases, had little effect (**Figure S5A, S5B**).

Crucially, astrocyte state reprogramming activity of gRNA-hit was abolished by introducing specific mismatches to the critical PAM-proximal motif GAGAG identified via ChIP-seq (**Figure 5C, right**). In particular, mutating bases 16-18 to T or C rather than the native ‘GAG’ resulted in the disruption of the critical motif and the complete blocking of cell state reprogramming, implying a removal of off-target effect. A possible explanation for this sequence-based activity difference is the reduced stability of gRNA secondary structures. However, we did not observe a strong relationship between minimum free energy (MFE) of simulated folded structures and reprogramming activity (**Figure S5C**). Therefore, we concluded that the GAGAG motif is a primary determinant of off-target site binding and bases within that motif are tolerant to DNA-substrate mismatches, but disrupting this motif blocks the promiscuous effects of this gRNA.

This screen provided information about cell state reprogramming via MAP2, but did not directly provide information regarding on-target activity of mutated variants at the FOXO4 promoter. Therefore, to explore the relationship between on and off-target activity, we selected the most and least active variants of gRNA-hit and control gRNA 23 (as measured by MAP2), and measured their on-target FOXO4 activation efficiency (**Figure 5D**). Interestingly, for gRNA-hit, the variants that most strongly upregulated MAP2, which contained mutations outside of the 5-bp critical PAM-proximal region of gRNA-hit, displayed decreased on-target FOXO4 activity. Introducing mismatches outside of the critical region may have decreased on-target site affinity but retained PAM-proximal base-driven off-target activity. However, the gRNA-hit variants with mismatches within these critical bases (and abolished MAP2 activity) displayed a surprising *increase* in on-target activity, activating

FOXO4 to ∼14-fold and ∼20-fold compared to non-targeting control. These findings suggest a potential relationship between on- and off-target activity, wherein widespread off-target activity may sequester dCas9 away from the on-target site, resulting in a lower effective on-target activation. This was further supported by gRNA-hit double mismatch data, where we observed that double mismatches introduced in positions 4 and 5 led to increased reprogramming activity (attributable by off-target effects) compared to double mismatches introduced in positions 1-2, where mismatches are almost completely tolerated and thus unlikely to impact binding or potential sequestration (**Figure 5E**).

### Off-target activation of PCDH7 by gRNA-hit contributes to astrocyte state reprogramming

We next set out to identify which genomic loci contribute to astrocyte reprogramming via off-target CRISPRa by gRNA-hit. We identified PCDH7 as a candidate active off-target due to its gRNA-hit ChIP-seq peak at the gene promoter (**Figure 3D**), PAM-dependent binding in our *in vitro* assay, and positive fold change in RNA-seq (**Figure S6A**). To test the contribution of PCDH7 to astrocyte state reprogramming, we directly targeted ^VP64^dSpCas9^VP64^ to the PCDH7 promoter, either alone or in the presence of gRNA-hit. As expected, gRNA-hit led to activation of FOXO4, but the PCDH7 gRNA did not (**Figure 6A**). In contrast, we observed a ∼60-fold activation of PCDH7 after delivery of only gRNA-hit, supporting off-target activity of gRNA-hit at the PCDH7 promoter. Directly targeting the PCDH7 promoter resulted in a 200-fold activation of PCDH7. Finally, we measured MAP2 expression as a state change proxy and found that activation of PCDH7 alone in absence of gRNA-hit nearly doubled MAP2 expression, supporting PCDH7 as contributing to astrocyte state change (**Figure 6A**). Together, these results demonstrate that individual functional off-targets of CRISPRa can be identified and validated experimentally. However, given the widespread genomic binding of CRISPRa with gRNA-hit, there are likely other loci, perhaps many, that contribute to state reprogramming.

**Figure 6:**
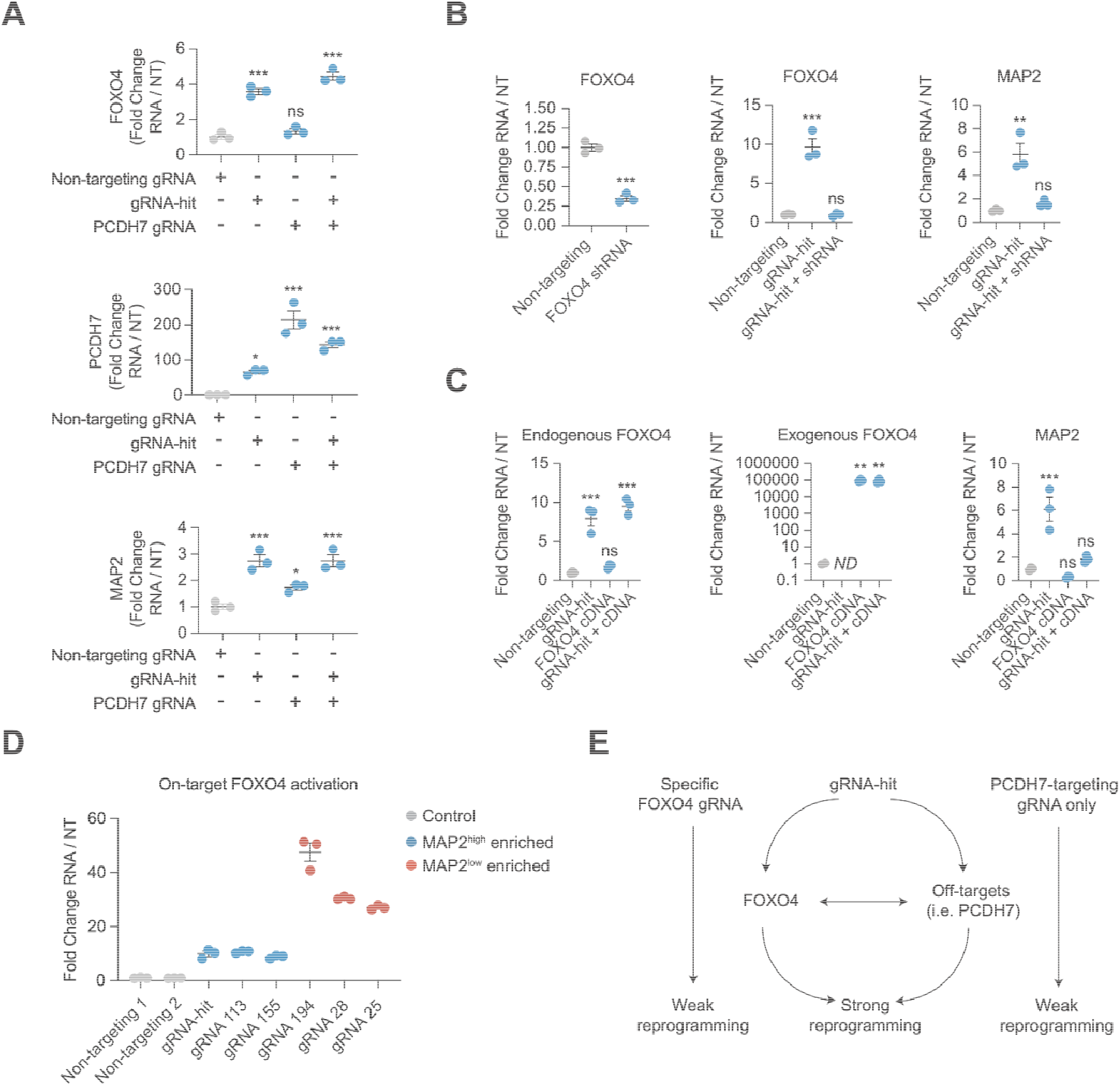
Simultaneous on-target and off-target activity contributes to transcriptional state reprogramming. A. RNA levels of FOXO4 (top), PCDH7 (middle), or MAP2 (bottom) after direct activation of PCDH7 alone or in combination with gRNA-hit. B. RNA levels of FOXO4 (left, middle), or MAP2 (right) after delivery of shRNA targeting FOXO4, alone or in combination with gRNA-hit. C. RNA levels of endogenous FOXO4 (left), exogenous codon-optimized FOXO4 (middle), or MAP2 (right) after delivery of FOXO4 cDNA, alone or in combination with gRNA-hit. In A-C, *p < 0.05, **p < 0.01, ***p < 0.001 by global one-way ANOVA with Dunnett’s post hoc test comparing all groups to non-targeting. Error bars represent SEM. D. RNA levels of FOXO4 after delivery of gRNAs identified in FOXO4 tiling screen to have a significant effect on MAP2 expression. gRNAs associated with astrocyte state reprogramming (enriched in MAP2^high^ cell populations) are shown in blue and display a similar level of FOXO4 activation as gRNA-hit. E. Model representing simultaneous functional on-target and off-target activity of gRNA-hit.

### Simultaneous on-target and off-target activity contributes to transcriptional state reprogramming

Our data support that gRNA-hit’s off-target activity drives astrocyte state change. We next sought to characterize the functional role, if any, of on-target activation of FOXO4 by gRNA-hit. First, to confirm that gRNA-hit reprograms in a Cas9-dependent manner, we expressed gRNA-hit in astrocytes without ^VP64^dSpCas9^VP64^ and found that no reprogramming was observed in absence of dCas9 (**Figure S6B**). Next, to untangle the effects of gRNA-hit’s on- and off-target activity, we delivered an shRNA targeting FOXO4 either alone or in the presence of gRNA-hit and measured astrocyte state reprogramming. First, we validated the FOXO4 shRNA by delivering it alone, which resulted in ∼70% knockdown of FOXO4 (**Figure 6B, left**).

Furthermore, when delivered alongside gRNA-hit, FOXO4 shRNA effectively knocked down FOXO4 expression back to WT levels (**Figure 6B, middle**). Critically, examination of MAP2 levels revealed that co-delivery of shRNA alongside gRNA-hit blocked the majority of state reprogramming, indicating that activation of FOXO4 is necessary for potent state reprogramming by gRNA-hit (**Figure 6B, right**). Therefore, we concluded that gRNA-hit reprograms astrocytes via simultaneous on-target activity at FOXO4 and off-target activity at sites including the PCDH7 promoter.

### On-target CRISPRa-FOXO4 astrocyte state reprogramming is dose-dependent

Finally, we explored the relationship between on-target FOXO4 activation and astrocyte state reprogramming. In cumulative testing of many FOXO4 gRNAs, we observed that despite activating FOXO4, some gRNAs did not display *reprogramming* activity (no change in MAP2 expression). TF expression level is tightly regulated, and precise dosage is often essential for cell differentiation and proper developmental patterning^90,91^. Furthermore, stronger TF activation by CRISPRa does not always correlate to a stronger downstream transcriptional impact^27^. Thus, we hypothesized that the contribution of CRISPRa-FOXO4 to state reprogramming is dose-dependent, such that activation of FOXO4 to a moderate level above baseline by gRNA-hit and other gRNAs is crucial for unlocking FOXO4’s contribution to astrocyte state reprogramming. To test this hypothesis, we tested FOXO4 activation at moderate and high levels alone and concurrently. To do so, we delivered a codon-optimized FOXO4 open reading frame (ORF) encoding the canonical FOXO4 transcript isoform either alone or in the presence of gRNA-hit and measured state reprogramming. CRISPRa with gRNA-hit activated endogenous FOXO4 to ∼8-fold of WT levels (**Figure 6C, left**). In contrast, delivery of FOXO4 ORF resulted in a significantly higher level of FOXO4, equating to >170-fold higher total FOXO4 RNA compared to gRNA-hit (**Figure 6C, middle**). We found that delivery of FOXO4 ORF did not alone result in state reprogramming. Separate delivery of each protein-coding isoform of FOXO4 without codon optimization also did not result in reprogramming (**Figure S6C**). Furthermore, when delivered together, FOXO4 ORF largely blocked state reprogramming potentiated by gRNA-hit (**Figure 6C, right**), implying that moderate FOXO4 activation could be crucial for astrocyte reprogramming. Finally, to confirm that careful titration of FOXO4 activation underlies FOXO4’s on-target contribution to cell reprogramming, we individually delivered the FOXO4 promoter-targeting gRNAs that alone resulted in moderate MAP2 increases in previous experiments (**Figure 2C**) and measured their on-target activation level. Indeed, the two gRNAs that emerged as positive hits from the promoter-tiling screen activated FOXO4 to a similar level as gRNA-hit, while three gRNAs that emerged from the screen as negative hits each activated FOXO4 to a significantly higher level, between 25-45-fold (**Figure 6D**). Thus, moderate activation is critical for unlocking FOXO4’s contribution to astrocyte state reprogramming.

## DISCUSSION

In this study, we identified a gRNA (“gRNA-hit”), selected to reprogram astrocyte cell state by targeting the promoter of FOXO4, with promiscuous and functional off-target activity. We show that ^VP64^dSpCas9^VP64^ guided by gRNA-hit drives the differential expression of thousands of genes, inducing a potent transcriptional state change in astrocytes. While this state can be modestly modulated by paired activation of other individual state-modifying TFs, the effects of gRNA-hit dominate the resulting phenotype. Astrocyte state is dynamic and heterogeneous, and cell atlases continue to uncover additional distinct transcriptional states in neural cells^92–94^. Researching astrocyte states will thus be important to advance understanding of reactive gliosis and neuroregeneration strategies like astrocyte-to-neuron reprogramming. It’s possible that widespread, functional, and sequence-informed off-target activity by gRNA-hit may result in activation of genes spanning pathways beyond those associated with glutamatergic neurons. Ultimately, the adequate fidelity of a transcriptional signature to the intended reprogrammed cell state will need to be determined on a case-by-case basis depending on the intended application.

CRISPRa is typically most effective when targeted slightly upstream of the TSS of its target gene ^95^. When we tested 198 FOXO4-targeting SpCas9 gRNAs within 1kb of the FOXO4 TSS, only gRNA-hit displayed a uniquely potent transcriptional state change. This unique effect prompted us to examine gRNA-hit’s off-target activity and we found by ChIP-seq that gRNA-hit binds reproducibly to thousands of sites across the human genome. The majority, but not all, of these sites contained a match for the 5bp PAM-proximal target sequence of gRNA-hit, followed by the NGG PAM. These sites spanned chromatin contexts. The majority fell in accessible chromatin but we observed >6,000 binding sites in inaccessible regions. However, these sites were enriched for promoters and intronic elements which are more likely to be accessible, highlighting the importance of chromatin state in determining off-target dCas9 binding.

ChIP-seq and similar methods including CUT&RUN^96^ and CUT&TAG^97^ provide valuable insight into dCas9 binding genome-wide but are labor-intensive and low-throughput. To gain insight into the mechanism of gRNA promiscuity in parallel to ChIP-seq, we employed high-throughput *in vitro* DNA binding arrays and gRNA-variant library screening approaches in cells. These complementary systems allow for systematic testing of DNA-target mismatches and gRNA-protospacer mismatches in high-throughput, revealing rules of mismatch tolerance. We observed that rG-dT mismatches were more tolerated than other base substitutions, especially in the PAM-proximal bases. We also observed that specific nucleotide positions within gRNAs differ in their mismatch tolerance, and that in certain cases introducing mismatches into the PAM-proximal critical region abolishes off-target activity but increases on-target activity. Previous reports have suggested that internal PAMs in gRNA seed regions modulate Cas9-DNA interaction and confer off-target binding in dCas9 systems^27,98^. Our study complements these reports by directly testing this hypothesis by editing bases within these motifs.

Overall, these data led us to a model of state reprogramming that depends on joint contributions of the on- and off-target activity of gRNA-hit, and a framework with which to interpret simultaneous on-target and promiscuous gRNA DNA-binding (**Figure 6E**). Individually, moderate on-target activity at the FOXO4 promoter has a relatively weak but significant effect on reprogramming and requires careful dose titration for effectiveness, such as with gRNA-hit, gRNA 113, and gRNA 155. In parallel, off-target effects, such as those contributed by PCDH7, individually contribute weak but significant reprogramming activity. By simultaneously leveraging functional on- and off-target activity, gRNA-hit potentiates particularly strong astrocyte state reprogramming, leading to potent transcriptional and phenotypic changes.

We note that gRNA-hit comes from a curated CRISPRa library^73^. It is likely that there are additional gRNAs within existing and widely-used libraries that display similar off-target characteristics as gRNA-hit. These problematic gRNAs could lead to false positives in screens that could be driving expression or cellular phenotypes in cryptic ways. Characterizing the gRNAs selected from these and other libraries with comprehensive and orthogonal methods such as ChIP-seq, DNA arrays, and gRNA-variant libraries will be critical to fully understand their specificity and mechanisms of action. As these datasets accumulate, integrative analyses can inform the extent of promiscuous activity in CRISPRi/a applications and the rules that govern off-target binding. This could ultimately lead to improved and more focused next-generation gRNA libraries.

Despite off-target activity typically being an unintended characteristic of CRISPR systems, this work demonstrates that in some instances, intended biological effects can be effectively coordinated by promiscuous gRNAs. This phenomenon may be particularly relevant for applications requiring broad transcriptional reprogramming, such as directing cell state changes. For example, we identify PCDH7 as a functional off-target site and demonstrate how simultaneous on- and off-target activity can synergize to produce a complex phenotype. Future work exploring multi-site genomic activity will likely be valuable for disentangling the relationship between on-target and off-target activity in these types of CRISPRa/i applications. Similarly, combinatorial perturbation is an inherently complex issue that can quickly scale to be experimentally unmanageable. However, promiscuous gRNAs could be advantageous in solving this complex problem. For example, as we further uncover the rules of on- and off-target binding, it may become possible to intentionally design gRNAs that target multiple sites specified by a short motif, such as GAGAG-NGG in the case of gRNA-hit, analogous to a synthetic TF that acts at many sites across the genome.

Finally, it is often assumed that the highest level of on-target activity is best for CRISPRa gRNA screens. In addition to our off-target characterization, these results demonstrate an example wherein the highest level of FOXO4 activation is not better, and instead establishes a nuanced requirement of modest FOXO4 expression for robust transcriptional state change. In addition to better understanding of off-target activity, future optimal CRISPRi/a gRNA library design may intentionally include a series of gRNAs for each target that elicit a distribution of effect sizes on target genes in order to fully capture these dose-responsive phenotypes.

## DATA AVAILABILITY

Data generated in high-throughput sequencing studies are available through NCBI GEO with accession numbers GSE311364, GSE311365, GSE311472 and GSE311503.

## SUPPLEMENTARY DATA

Supplementary data are available at NAR online.

## ACKNOWLEDGEMENTS

We thank all members of the Gersbach laboratory, Ruhi Rai, and Alejandro Barrera for technical assistance and helpful discussions. Figure 3B was created using https://sankeymatic.com/.

Author contributions: SJR, WZ, RG, and CAG designed experiments. SJR, WZ, SEM, DH and NS performed the experiments. WZ performed the in vitro dCas9 binding experiments. SJR analyzed experimental data.

GEC, RG, and CAG supervised the study. SJR and CAG wrote the manuscript with contributions by all authors.

## REFERENCES

1. McCutcheon, S. R., Rohm, D., Iglesias, N. & Gersbach, C. A. Epigenome editing technologies for discovery and medicine. Nat. Biotechnol. 42, 1199–1217 (2024).

2. Dominguez, A. A., Lim, W. A. & Qi, L. S. Beyond editing: repurposing CRISPR-Cas9 for precision genome regulation and interrogation. Nat. Rev. Mol. Cell Biol. 17, 5–15 (2016).

3. Kampmann, M. CRISPRi and CRISPRa Screens in Mammalian Cells for Precision Biology and Medicine. ACS Chem. Biol. 13, 406–416 (2018).

4. Klann, T. S., Black, J. B. & Gersbach, C. A. CRISPR-based methods for high-throughput annotation of regulatory DNA. Curr. Opin. Biotechnol. 52, 32–41 (2018).

5. Butterfield, G. L., Reisman, S. J., Iglesias, N. & Gersbach, C. A. Gene regulation technologies for gene and cell therapy. Mol. Ther. J. Am. Soc. Gene Ther. 33, 2104–2122 (2025).

6. Cullot, G. et al. CRISPR-Cas9 genome editing induces megabase-scale chromosomal truncations. Nat. Commun. 10, 1136 (2019).

7. Weltner, J. et al. Human pluripotent reprogramming with CRISPR activators. Nat. Commun. 9, 2643 (2018).

8. Kwon, J. B., Vankara, A., Ettyreddy, A. R., Bohning, J. D. & Gersbach, C. A. Myogenic Progenitor Cell Lineage Specification by CRISPR/Cas9-Based Transcriptional Activators. Stem Cell Rep. 14, 755–769 (2020).

9. Liu, Y. et al. CRISPR Activation Screens Systematically Identify Factors that Drive Neuronal Fate and Reprogramming. Cell Stem Cell 23, 758–771.e8 (2018).

10. Black, J. B. et al. Master Regulators and Cofactors of Human Neuronal Cell Fate Specification Identified by CRISPR Gene Activation Screens. Cell Rep. 33, 108460 (2020).

11. Chakraborty, S. et al. A CRISPR/Cas9-Based System for Reprogramming Cell Lineage Specification. Stem Cell Rep. 3, 940–947 (2014).

12. Black, J. B. et al. Targeted Epigenetic Remodeling of Endogenous Loci by CRISPR/Cas9-Based Transcriptional Activators Directly Converts Fibroblasts to Neuronal Cells. Cell Stem Cell 19, 406–414 (2016).

13. Chavez, A. et al. Highly efficient Cas9-mediated transcriptional programming. Nat. Methods 12, 326–328 (2015).

14. Shakirova, K. M., Ovchinnikova, V. Y. & Dashinimaev, E. B. Cell Reprogramming With CRISPR/Cas9 Based Transcriptional Regulation Systems. Front. Bioeng. Biotechnol. 8, 882 (2020).

15. Datlinger, P. et al. Pooled CRISPR screening with single-cell transcriptome readout. Nat. Methods 14, 297–301 (2017).

16. Bock, C. et al. High-content CRISPR screening. Nat. Rev. Methods Primer 2, 9 (2022).

17. Rohm, D. et al. Activation of the imprinted Prader-Willi syndrome locus by CRISPR-based epigenome editing. Cell Genomics 5, 100770 (2025).

18. Gasperini, M. et al. A Genome-wide Framework for Mapping Gene Regulation via Cellular Genetic Screens. Cell 176, 1516 (2019).

19. Replogle, J. M. et al. Mapping information-rich genotype-phenotype landscapes with genome-scale Perturb-seq. Cell 185, 2559–2575.e28 (2022).

20. Tsai, S. Q. et al. CIRCLE-seq: a highly sensitive in vitro screen for genome-wide CRISPR-Cas9 nuclease off-targets. Nat. Methods 14, 607–614 (2017).

21. Tsai, S. Q. et al. GUIDE-seq enables genome-wide profiling of off-target cleavage by CRISPR-Cas nucleases. Nat. Biotechnol. 33, 187–197 (2015).

22. Kim, D. et al. Digenome-seq: genome-wide profiling of CRISPR-Cas9 off-target effects in human cells. Nat. Methods 12, 237–243, 1 p following 243 (2015).

23. Wienert, B. et al. Unbiased detection of CRISPR off-targets in vivo using DISCOVER-Seq. Science 364, 286–289 (2019).

24. Perez-Pinera, P. et al. RNA-guided gene activation by CRISPR-Cas9–based transcription factors. Nat. Methods 10, 973–976 (2013).

25. Polstein, L. R. et al. Genome-wide specificity of DNA binding, gene regulation, and chromatin remodeling by TALE- and CRISPR/Cas9-based transcriptional activators. Genome Res. 25, 1158–1169 (2015).

26. Gemberling, M. P. et al. Transgenic mice for in vivo epigenome editing with CRISPR-based systems. Nat. Methods 18, 965–974 (2021).

27. Southard, K. M. et al. Comprehensive transcription factor perturbations recapitulate fibroblast transcriptional states. Nat. Genet. 57, 2323–2334 (2025).

28. Kuscu, C., Arslan, S., Singh, R., Thorpe, J. & Adli, M. Genome-wide analysis reveals characteristics of off-target sites bound by the Cas9 endonuclease. Nat. Biotechnol. 32, 677–683 (2014).

29. Thakore, P. I. et al. Highly specific epigenome editing by CRISPR-Cas9 repressors for silencing of distal regulatory elements. Nat. Methods 12, 1143–1149 (2015).

30. Rohatgi, N. et al. Seed sequences mediate off-target activity in the CRISPR-interference system. Cell Genomics 4, 100693 (2024).

31. Fu, Y. et al. High-frequency off-target mutagenesis induced by CRISPR-Cas nucleases in human cells. Nat. Biotechnol. 31, 822–826 (2013).

32. Chen, B. et al. Dynamic imaging of genomic loci in living human cells by an optimized CRISPR/Cas system. Cell 155, 1479–1491 (2013).

33. McCutcheon, S. R. et al. Transcriptional and epigenetic regulators of human CD8+ T cell function identified through orthogonal CRISPR screens. Nat. Genet. 55, 2211–2223 (2023).

34. Gordon, M. G. et al. lentiMPRA and MPRAflow for high-throughput functional characterization of gene regulatory elements. Nat. Protoc. 15, 2387–2412 (2020).

35. Bolger, A. M., Lohse, M. & Usadel, B. Trimmomatic: a flexible trimmer for Illumina sequence data. Bioinforma. Oxf. Engl. 30, 2114–2120 (2014).

36. Dobin, A. et al. STAR: ultrafast universal RNA-seq aligner. Bioinforma. Oxf. Engl. 29, 15–21 (2013).

37. Liao, Y., Smyth, G. K. & Shi, W. The Subread aligner: fast, accurate and scalable read mapping by seed-and-vote. Nucleic Acids Res. 41, e108 (2013).

38. Love, M. I., Huber, W. & Anders, S. Moderated estimation of fold change and dispersion for RNA-seq data with DESeq2. Genome Biol. 15, 550 (2014).

39. Chen, E. Y. et al. Enrichr: interactive and collaborative HTML5 gene list enrichment analysis tool. BMC Bioinformatics 14, 128 (2013).

40. Reisman, S. J. et al. Comprehensive profiling of transcription factors for reprogramming human astrocytes to neuronal cells through endogenous CRISPR-based gene activation. Preprint at 10.1101/2025.10.11.681828 (2025).

41. Schmidt, H. et al. Genome-wide CRISPR guide RNA design and specificity analysis with GuideScan2. Genome Biol. 26, 41 (2025).

42. Lorenz, R. et al. ViennaRNA Package 2.0. Algorithms Mol. Biol. AMB 6, 26 (2011).

43. Langmead, B. & Salzberg, S. L. Fast gapped-read alignment with Bowtie 2. Nat. Methods 9, 357–359 (2012).

44. Hao, Y. et al. Integrated analysis of multimodal single-cell data. Cell 184, 3573–3587.e29 (2021).

45. Hafemeister, C. & Satija, R. Normalization and variance stabilization of single-cell RNA-seq data using regularized negative binomial regression. Genome Biol. 20, 296 (2019).

46. Liu, S. et al. Characterization and bioinformatic filtering of ambient gRNAs in single-cell CRISPR screens using CLEANSER. Cell Genomics 5, 100766 (2025).

47. Finak, G. et al. MAST: a flexible statistical framework for assessing transcriptional changes and characterizing heterogeneity in single-cell RNA sequencing data. Genome Biol. 16, 278 (2015).

48. Li, H. et al. The Sequence Alignment/Map format and SAMtools. Bioinformatics 25, 2078–2079 (2009).

49. Zhang, Y. et al. Model-based analysis of ChIP-Seq (MACS). Genome Biol. 9, R137 (2008).

50. Li, Q., Brown, J. B., Huang, H. & Bickel, P. J. Measuring reproducibility of high-throughput experiments. Ann. Appl. Stat. 5, (2011).

51. Quinlan, A. R. BEDTools: The Swiss-Army Tool for Genome Feature Analysis. Curr. Protoc. Bioinforma. 47, 11.12.1-34 (2014).

52. Ramírez, F., Dündar, F., Diehl, S., Grüning, B. A. & Manke, T. deepTools: a flexible platform for exploring deep-sequencing data. Nucleic Acids Res. 42, W187–191 (2014).

53. Robinson, J. T. et al. Integrative genomics viewer. Nat. Biotechnol. 29, 24–26 (2011).

54. Heinz, S. et al. Simple combinations of lineage-determining transcription factors prime cis-regulatory elements required for macrophage and B cell identities. Mol. Cell 38, 576–589 (2010).

55. Vilchez, D. et al. FOXO4 is necessary for neural differentiation of human embryonic stem cells. Aging Cell 12, 518–522 (2013).

56. Joung, J. et al. A transcription factor atlas of directed differentiation. Cell 186, 209–229.e26 (2023).

57. Liddelow, S. A. & Barres, B. A. Reactive Astrocytes: Production, Function, and Therapeutic Potential. Immunity 46, 957–967 (2017).

58. Alizadeh, S. D. et al. Reprogramming of astrocytes to neuronal-like cells in spinal cord injury: a systematic review. Spinal Cord 62, 133–142 (2024).

59. Gao, L. et al. Direct Generation of Human Neuronal Cells from Adult Astrocytes by Small Molecules. Stem Cell Rep. 8, 538–547 (2017).

60. Herrero-Navarro, Á. et al. Astrocytes and neurons share region-specific transcriptional signatures that confer regional identity to neuronal reprogramming. Sci. Adv. 7, eabe8978 (2021).

61. Lentini, C. et al. Reprogramming reactive glia into interneurons reduces chronic seizure activity in a mouse model of mesial temporal lobe epilepsy. Cell Stem Cell 28, 2104–2121.e10 (2021).

62. Rivetti di Val Cervo, P. et al. Induction of functional dopamine neurons from human astrocytes in vitro and mouse astrocytes in a Parkinson’s disease model. Nat. Biotechnol. 35, 444–452 (2017).

63. Vasan, L., Park, E., David, L. A., Fleming, T. & Schuurmans, C. Direct Neuronal Reprogramming: Bridging the Gap Between Basic Science and Clinical Application. Front. Cell Dev. Biol. 9, 681087 (2021).

64. Wu, Z. et al. Gene therapy conversion of striatal astrocytes into GABAergic neurons in mouse models of Huntington’s disease. Nat. Commun. 11, 1105 (2020).

65. Yin, J.-C. et al. Chemical Conversion of Human Fetal Astrocytes into Neurons through Modulation of Multiple Signaling Pathways. Stem Cell Rep. 12, 488–501 (2019).

66. Zhang, L. et al. Small Molecules Efficiently Reprogram Human Astroglial Cells into Functional Neurons. Cell Stem Cell 17, 735–747 (2015).

67. Zhang, Y. et al. A single factor elicits multilineage reprogramming of astrocytes in the adult mouse striatum. Proc. Natl. Acad. Sci. U. S. A. 119, e2107339119 (2022).

68. Dehmelt, L. & Halpain, S. The MAP2/Tau family of microtubule-associated proteins. Genome Biol. 6, 204 (2005).

69. Vilchez, D. et al. FOXO4 is necessary for neural differentiation of human embryonic stem cells. Aging Cell 12, 518–522 (2013).

70. Baar, M. P. et al. Targeted Apoptosis of Senescent Cells Restores Tissue Homeostasis in Response to Chemotoxicity and Aging. Cell 169, 132–147.e16 (2017).

71. Fremeau, R. T. et al. The expression of vesicular glutamate transporters defines two classes of excitatory synapse. Neuron 31, 247–260 (2001).

72. Dixit, A. et al. Perturb-Seq: Dissecting Molecular Circuits with Scalable Single-Cell RNA Profiling of Pooled Genetic Screens. Cell 167, 1853–1866.e17 (2016).

73. Sanson, K. R. et al. Optimized libraries for CRISPR-Cas9 genetic screens with multiple modalities. Nat. Commun. 9, 5416 (2018).

74. Perez, A. R. et al. GuideScan software for improved single and paired CRISPR guide RNA design. Nat. Biotechnol. 35, 347–349 (2017).

75. Guo, C., Ma, X., Gao, F. & Guo, Y. Off-target effects in CRISPR/Cas9 gene editing. Front. Bioeng. Biotechnol. 11, 1143157 (2023).

76. Berger, M. F. et al. Compact, universal DNA microarrays to comprehensively determine transcription-factor binding site specificities. Nat. Biotechnol. 24, 1429–1435 (2006).

77. Berger, M. F. & Bulyk, M. L. Protein binding microarrays (PBMs) for rapid, high-throughput characterization of the sequence specificities of DNA binding proteins. Methods Mol. Biol. Clifton NJ 338, 245–260 (2006).

78. Wu, X. et al. Genome-wide binding of the CRISPR endonuclease Cas9 in mammalian cells. Nat. Biotechnol. 32, 670–676 (2014).

79. Pattanayak, V. et al. High-throughput profiling of off-target DNA cleavage reveals RNA-programmed Cas9 nuclease specificity. Nat. Biotechnol. 31, 839–843 (2013).

80. Jinek, M. et al. A programmable dual-RNA-guided DNA endonuclease in adaptive bacterial immunity. Science 337, 816–821 (2012).

81. Hsu, P. D. et al. DNA targeting specificity of RNA-guided Cas9 nucleases. Nat. Biotechnol. 31, 827–832 (2013).

82. Cong, L. et al. Multiplex genome engineering using CRISPR/Cas systems. Science 339, 819–823 (2013).

83. Zhu, W. et al. dCas9 binding specificity measured on high-density DNA arrays reveals insights into guide inefficiencies and off-target binding. Prep.

84. Kimsey, I. J., Petzold, K., Sathyamoorthy, B., Stein, Z. W. & Al-Hashimi, H. M. Visualizing transient Watson-Crick-like mispairs in DNA and RNA duplexes. Nature 519, 315–320 (2015).

85. Pacesa, M. et al. Structural basis for Cas9 off-target activity. Cell 185, 4067–4081.e21 (2022).

86. Anderson, E. M. et al. Systematic analysis of CRISPR-Cas9 mismatch tolerance reveals low levels of off-target activity. J. Biotechnol. 211, 56–65 (2015).

87. Pan, X. et al. Massively targeted evaluation of therapeutic CRISPR off-targets in cells. Nat. Commun. 13, 4049 (2022).

88. Boyle, E. A. et al. High-throughput biochemical profiling reveals sequence determinants of dCas9 off-target binding and unbinding. Proc. Natl. Acad. Sci. U. S. A. 114, 5461–5466 (2017).

89. Jost, M. et al. Titrating gene expression using libraries of systematically attenuated CRISPR guide RNAs. Nat. Biotechnol. 38, 355–364 (2020).

90. Ni, Z., Zhou, X.-Y., Aslam, S. & Niu, D.-K. Characterization of Human Dosage-Sensitive Transcription Factor Genes. Front. Genet. 10, 1208 (2019).

91. Lundgren, M. et al. Transcription factor dosage affects changes in higher order chromatin structure associated with activation of a heterochromatic gene. Cell 103, 733–743 (2000).

92. Rafelski, S. M. & Theriot, J. A. Establishing a conceptual framework for holistic cell states and state transitions. Cell 187, 2633–2651 (2024).

93. Bocchi, R. et al. Astrocyte heterogeneity reveals region-specific astrogenesis in the white matter. Nat. Neurosci. 28, 457–469 (2025).

94. Chen, X. et al. A brain cell atlas integrating single-cell transcriptomes across human brain regions. Nat. Med. 30, 2679–2691 (2024).

95. Gilbert, L. A. et al. Genome-Scale CRISPR-Mediated Control of Gene Repression and Activation. Cell 159, 647–661 (2014).

96. Skene, P. J. & Henikoff, S. An efficient targeted nuclease strategy for high-resolution mapping of DNA binding sites. eLife 6, e21856 (2017).

97. Kaya-Okur, H. S. et al. CUT&Tag for efficient epigenomic profiling of small samples and single cells. Nat. Commun. 10, 1930 (2019).

98. Corsi, G. I. et al. CRISPR/Cas9 gRNA activity depends on free energy changes and on the target PAM context. Nat. Commun. 13, 3006 (2022).

