## Supplementary Figures for "Mismatch tolerance of a gRNA for CRISPR-based gene activation confers broad activity critical for cell reprogramming"

Figure S1: Details of astrocyte transcriptional reprogramming by gRNA-hit (Related to Figure 1).

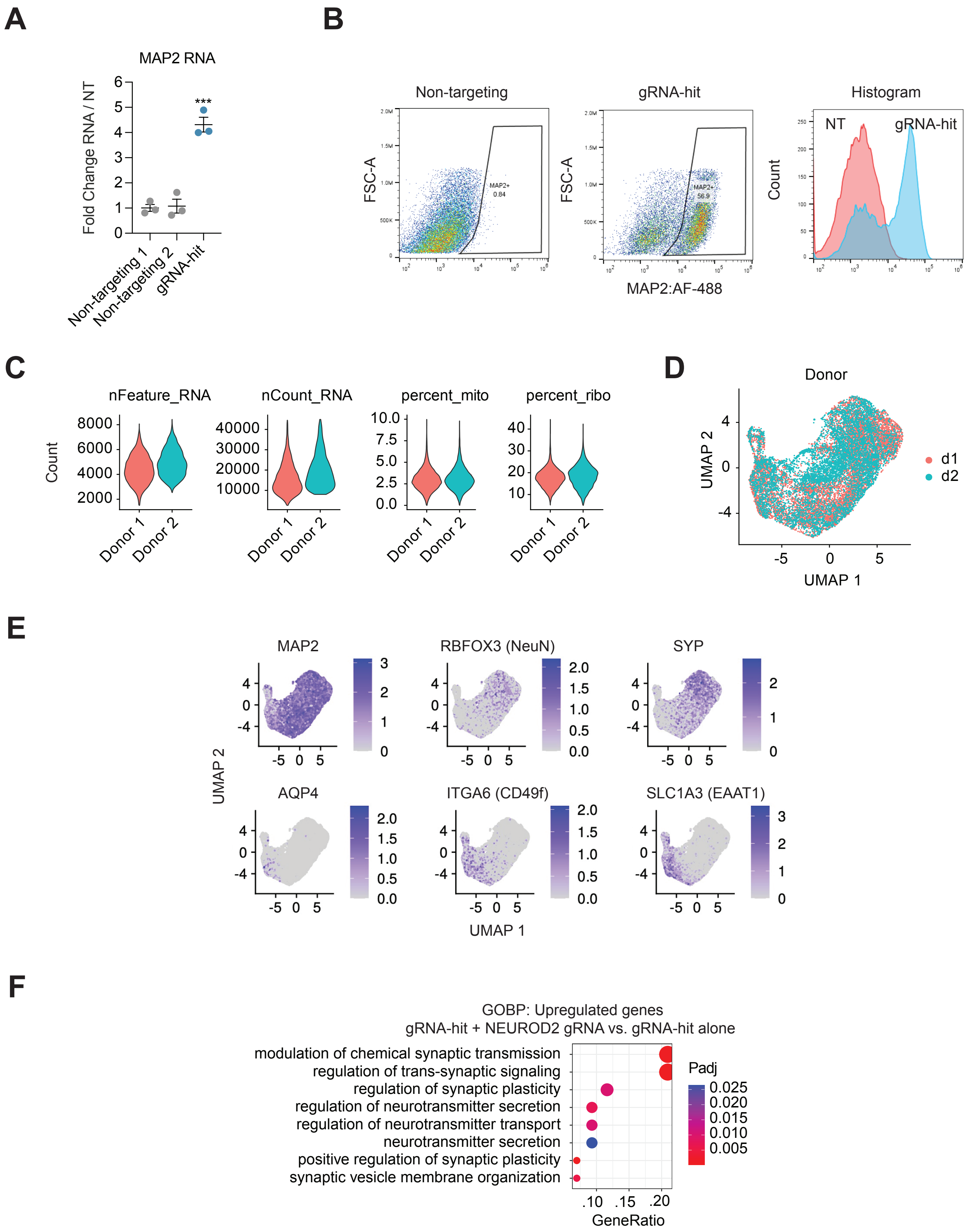

**Figure S1: Details of astrocyte transcriptional reprogramming by gRNA-hit (Related to Figure 1).**

A. MAP2 RNA expression after CRISPRa-FOXO4 with gRNA-hit, 10 days post-transduction. ***p < 0.001 by global one-way ANOVA with Dunnett's post hoc test comparing all groups to non-targeting 1; error bars represent SEM.
B. Representative FACS scatter displaying MAP2 protein expression after CRISPRa-FOXO4 with gRNA-hit.
C. Cell QC metrics for each astrocyte donor profiled in the perturb-seq dataset.
D. UMAP embedding colored by donor.
E. Feature plots displaying count distribution of neuron marker genes (top) and astrocyte marker genes (bottom) in UMAP distribution.
F. Top enriched biological processes for upregulated DE genes detected in perturb-seq. Contrast: gRNA-hit + NEUROD2 gRNA vs. gRNA-hit alone. Statistical significance of term enrichment was determined using a two-tailed Fisher's exact test followed by Benjamini–Hochberg correction.

Figure S2: Differential expression after delivery of FOXO4-targeting gRNAs (Related to Figure 2).

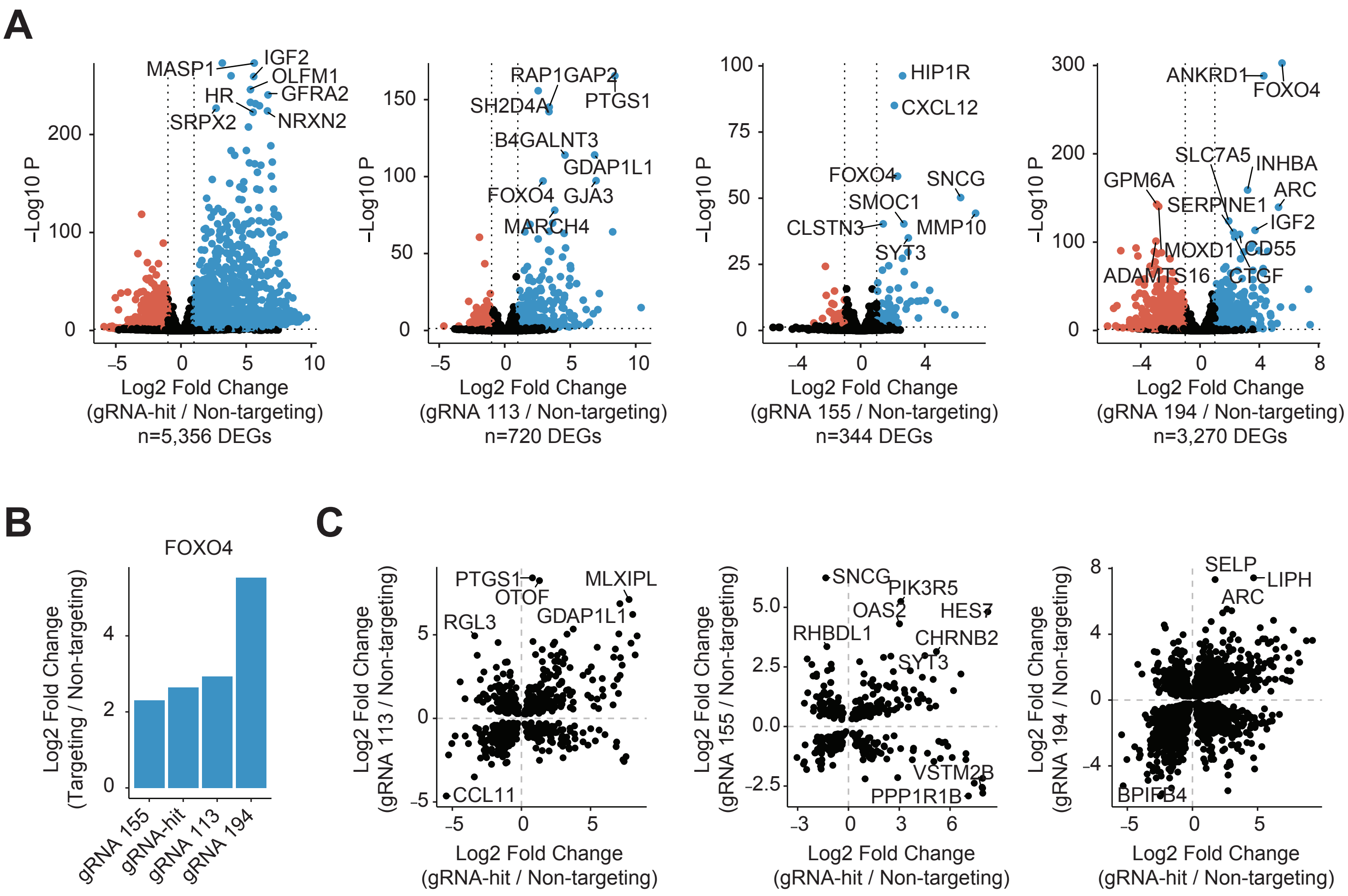

**Figure S2: Differential expression after delivery of FOXO4-targeting gRNAs (Related to Figure 2).**

A. Volcano plots of DE genes after delivery of FOXO4-targeting gRNAs. DEGs are defined by Padj < 0.01 by differential DESeq2 analysis.
B. On-target FOXO4 activation levels after delivery of FOXO4-targeting gRNAs.
C. Scatterplots comparing fold changes of shared DE genes after delivery of FOXO4-targeting gRNAs.

Figure S3: Details of $^{VP64}$dSpCas9$^{VP64}$ ChIP-seq (Related to Figure 3).

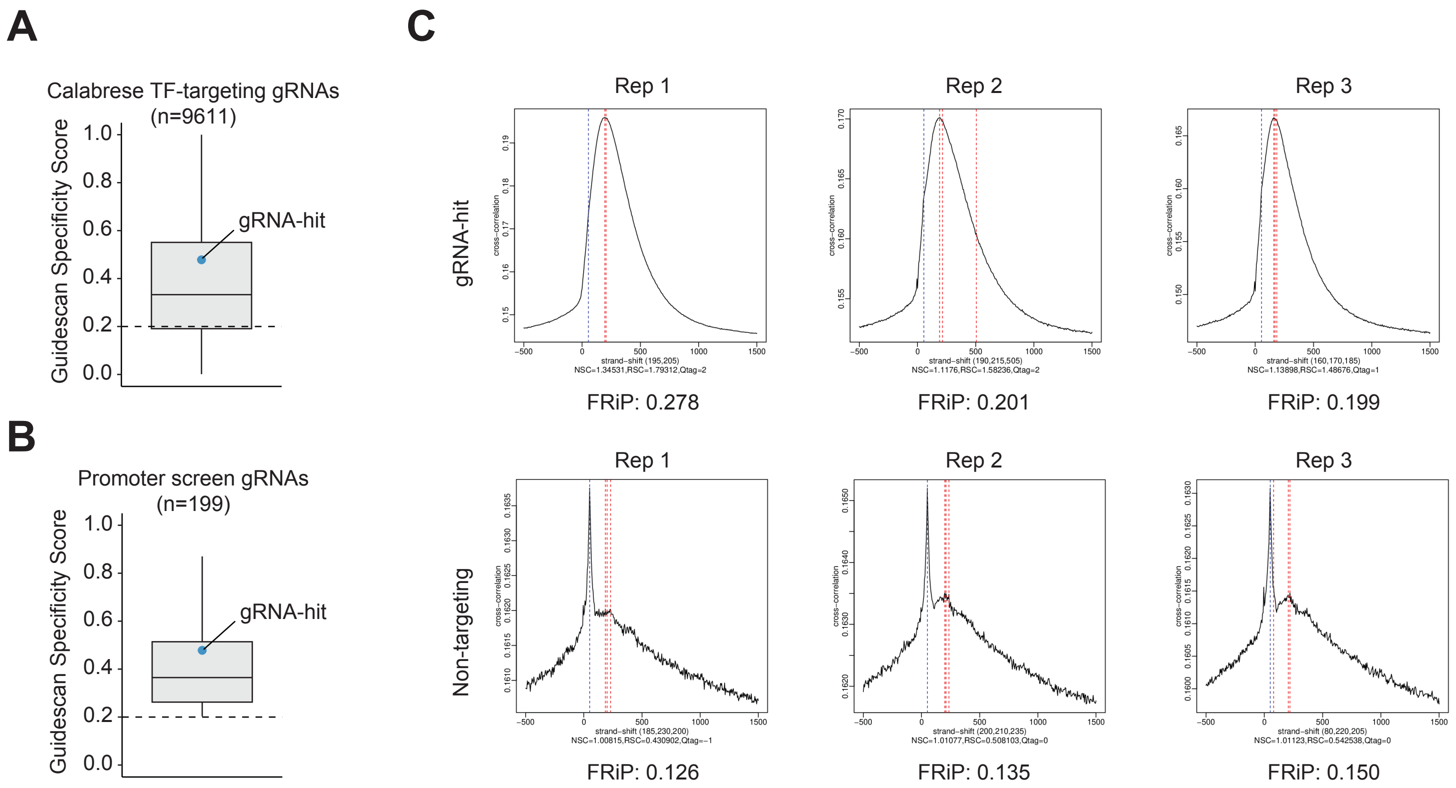

**Figure S3: Details of $^{VP64}$dSpCas9$^{VP64}$ ChIP-seq (Related to Figure 3).**

A. Guidescan specificity scores for all Calabrese gRNAs [PMID: 30575746]. The blue point denotes gRNA-hit.
B. Guidescan specificity scores for all FOXO4 promoter screen gRNAs. The blue point denotes gRNA-hit.
C. ENCODE ChIP-seq pipeline QC metrics, strand cross-correlation (SSC) and Fraction reads in peak (FRiP).

Figure S4: Details of DNA target mismatch tolerance (Related to Figure 4).

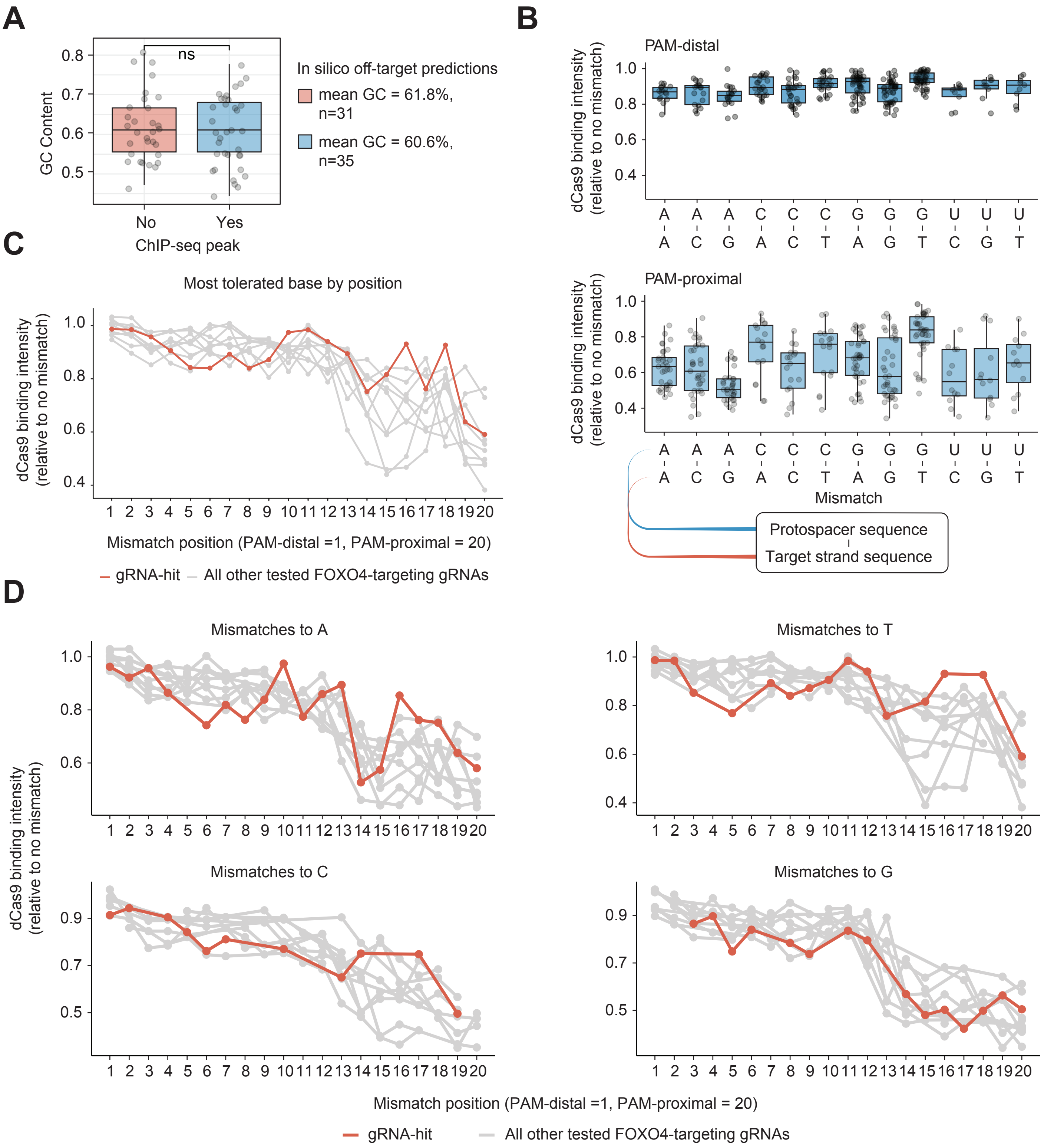

**Figure S4: Details of DNA target mismatch tolerance (Related to Figure 4).**

A. GC content of in silico gRNA-hit off-target sites, separated by whether they fell in a ChIP-seq peak. ns > 0.05 by two-sided Wilcoxon rank-sum test.
B. Protospacer-DNA-target mismatch tolerance. Binding is relative to gRNA-matched on-target binding. PAM-distal and PAM-proximal bases were divided evenly, 10 bases per category.
C. Comparison of binding intensities of most-tolerated mismatch by position and gRNA. gRNA-hit is shown in red, 9 other FOXO4-targeting gRNAs are shown in gray.
D. Comparison of binding intensities of mismatches to each new base in DNA-target. gRNA-hit is shown in red, 9 other FOXO4-targeting gRNAs are shown in gray. Missing points correspond to where the plotted base was already present at that position in that gRNA's DNA-target.

Figure S5: Details of gRNA variant screening (Related to Figure 5).

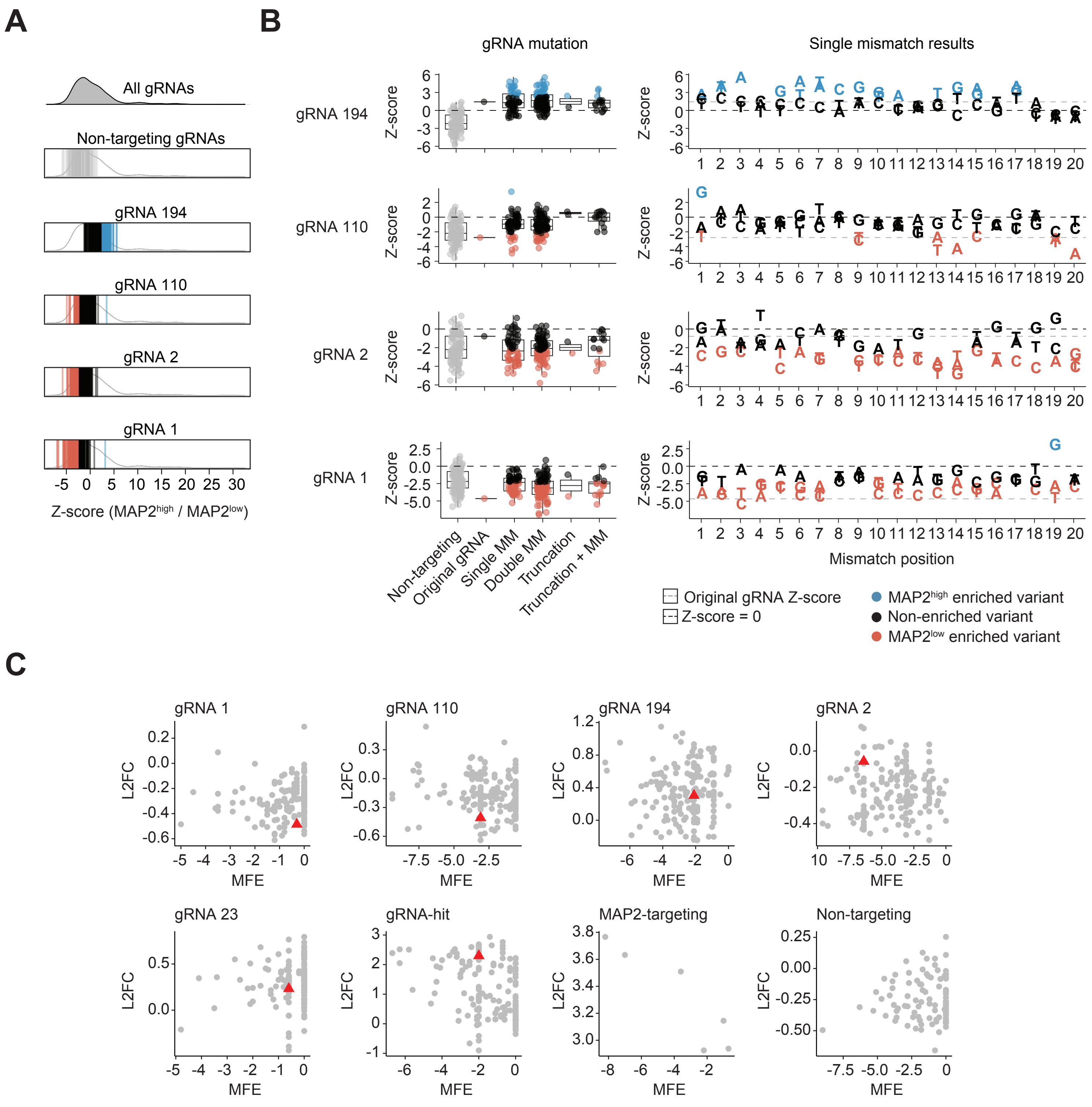

**Figure S5: Details of gRNA variant screening (Related to Figure 5).**

A. Screen DESeq2 Z-score of gRNA variants. Enriched variants (Padj < 0.01) were defined using a paired two-tailed DESeq2 test with Benjamini–Hochberg correction. The overall distribution is slightly left-shifted due to strong positive fold change of gRNA-hit variants.
B. gRNA variant screen DESeq2 Z-score by category (left) and single mismatch variants by base position (right) for additional control gRNAs (gRNA 194, gRNA 110, gRNA 2, gRNA 1). Non-mutated gRNA Z-score is represented by a gray dashed line, Z-score = 0 is represented by a black dashed line, and plotted bases represent new DNA bases encoding the gRNA at each position.
C. Scatterplots comparing predicted minimum free energy of gRNA variant secondary structures to screen fold changes. Non-mutated gRNAs are represented by a red triangle, all other variants are represented by gray circles.

Figure S6: Details of on-target and off-target activity of gRNA-hit (Related to Figure 6).

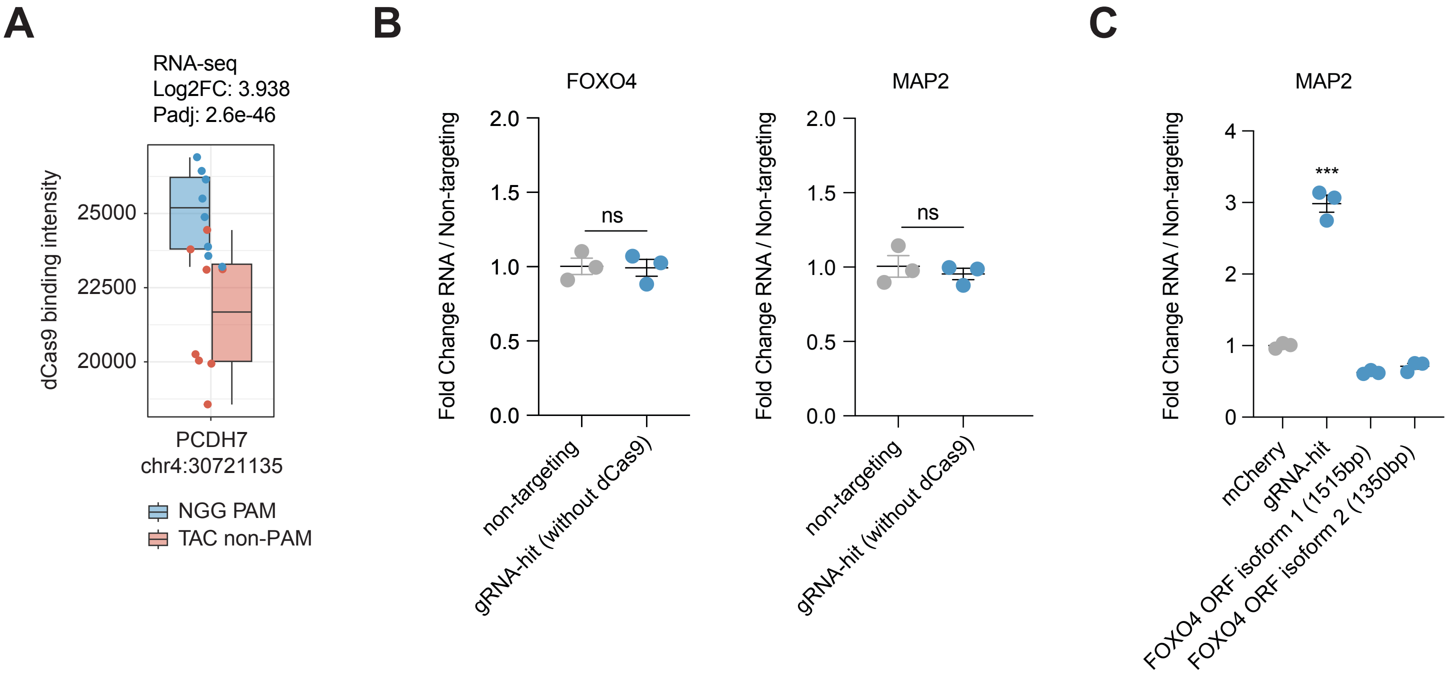

**Figure S6: Details of on-target and off-target activity of gRNA-hit (Related to Figure 6).**

A. Binding of dCas9 to PCDH7 promoter sequence (predicted off-target) in *in vitro* binding assay (Figure 4) when guided by gRNA-hit.
B. RNA levels of FOXO4 (left) and MAP2 (right) after delivery of gRNA-hit in the absence of $^{VP64}$dSpCas9$^{VP64}$.
C. RNA levels of MAP2 after delivery of ORFs encoding the 2 canonical FOXO4 isoforms or gRNA-hit. In B-C, ns = $p > 0.05$, ***$p < 0.001$ by global one-way ANOVA with Dunnett's post hoc test comparing all groups to non-targeting. Error bars represent SEM.
